# The killifish germline regulates longevity and somatic repair in a sex-specific manner

**DOI:** 10.1101/2023.12.18.572041

**Authors:** Eitan Moses, Tehila Atlan, Xue Sun, Roman Franek, Atif Siddiqui, Georgi K. Marinov, Sagiv Shifman, David M. Zucker, Adi Oron-Gottesman, William J. Greenleaf, Ehud Cohen, Oren Ram, Itamar Harel

## Abstract

Classical evolutionary theories propose tradeoffs between reproduction, damage repair, and lifespan. However, the specific role of the germline in shaping vertebrate aging remains largely unknown. Here, we use the turquoise killifish (*N. furzeri*) to genetically arrest germline development at discrete stages, and examine how different modes of infertility impact life-history. We first construct a comprehensive single-cell gonadal atlas, providing cell-type-specific markers for downstream phenotypic analysis. Next, we show that germline depletion - but not arresting germline differentiation - enhances damage repair in female killifish. Conversely, germline-depleted males instead showed an extension in lifespan and rejuvenated metabolic functions. Through further transcriptomic analysis, we highlight enrichment of pro-longevity pathways and genes in germline-depleted male killifish and demonstrate functional conservation of how these factors may regulate longevity in germline-depleted *C. elegans*. Our results therefore demonstrate that different germline manipulation paradigms can yield pronounced sexually dimorphic phenotypes, implying alternative responses to classical evolutionary tradeoffs.

## Introduction

For many species age at maturity is proportional to adult lifespan^1,2^. This correlation has sparked several evolutionary theories proposing a functional trade-off between life-history traits^3,4^. One of the leading concepts, the ‘disposable soma’ theory, suggests that there is a tradeoff between reproduction, growth, damage repair (e.g. DNA damage), and organismal lifespan^5,6^. Accordingly, a recent study in zebrafish has proposed that germline repair itself can have a direct tradeoff with somatic regeneration^7^. These findings suggest that competing demands can limit the resources that are available for repairing somatic damage, which could ultimately lead to damage accumulation and may affect lifespan.

Alternatively, seminal experiments in *C. elegans* have demonstrated that ablating the germline can produce remarkable lifespan extension^8^, but only when the empty somatic gonad remains intact (i.e. not via full castration). These findings suggest that the longevity phenotype is not just a consequence of a trade-off in resources, but rather depends on endocrine signals mediated by the gonad; either pro-longevity signals released by an empty gonad or loss of pro-aging signals released from the intact gonad. Downstream, enhanced pro-longevity pathways were detected in germline-ablated worms, including enhanced proteostasis and lipid metabolism^9^. Intriguing correlative data exists in humans, as well as several farm and laboratory animals, whereby removal of both the somatic gonad and the germline through castration (e.g. in Korean eunuchs) can have a profound effect on longevity^10–18^. Thus, additional experimental evidence is required to determine how the germline itself can impact vertebrate lifespan, and to elucidate the relative contribution of each of the seemingly opposing theories described above^19–22^.

Unlike worms, vertebrate reproduction is regulated by the vertebrate-specific hypothalamic– pituitary–gonadal (HPG) axis. Therefore, paradigms cannot be simply extrapolated from studies in *C. elegans*. In addition, most vertebrates are segregated into true females and males, thereby introducing the possibility of investigating sex-specific effects. Exploring the described hypotheses in a vertebrate model may therefore provide insights concerning sexual dimorphism in lifespan regulation^23^. However, moving away from invertebrates to a vertebrate model introduces new challenges, one of which is the relatively long lifespan of classical vertebrate models that hampers large-scale genetic interrogations. Therefore, the development of alternative experimental strategies is required.

As the most diverse group of vertebrates, fish species exhibit a vast range of life-histories^24,25^. For example, the long-lived rockfish and Greenland sharks mature after decades (and even centuries)^26,27^. At the other extreme, the naturally short-lived African turquoise killifish (*Nothobranchius furzeri*) can undergo sexual maturation as fast as 14 days^28^, and exhibits a median lifespan of only ∼4-6 months^29^. Accordingly, the naturally compressed lifespan of the killifish is 6-10-fold shorter than the lifespan of mice and zebrafish, respectively. Thus, providing a unique opportunity for rapid exploration of vertebrate physiology. The turquoise killifish has vertebrate-conserved organs and systems, including the HPG reproductive axis, and a recently developed state-of-the-art genome engineering platform^30–38^. However, the regulatory mechanisms of reproduction and gonadal development in killifish are largely unknown.

Most genetic studies of fish reproduction have used the zebrafish (*Danio rerio*) and medaka (*Oryzias latipes*) models, but these exhibit unique characteristics. For example, both of these fish depend on the presence of the germline to develop into females (i.e. sterile fish undergo an all-male sex-reversal)^39,40^, and zebrafish lack an XY-based sexual determination system^41^. Thus, although recent studies have begun to characterize fish gonads at the single cell level^42,43^, it is currently unknown which patterns are conserved in killifish.

Here, we leverage the turquoise killifish to dissect the link between reproduction and vertebrate lifespan. To this end, we first generate a gonadal cell-type atlas at a single cell resolution, in both male and female fish. This detailed resource allows us to identify primary gonadal cell-types and provide molecular markers for downstream phenotypic analysis. We then generate a panel of genetic models to perturb the reproductive state in killifish, and measure the tradeoff with key physiological properties, including somatic growth and damage repair. Thus, allowing us to dissect the systemic effects of distinct paradigms of germline manipulation in a vertebrate model.

Our results indicate that only depleting the germline, and not blocking germline differentiation, can enhance female-specific somatic repair. To further explore these sex differences, we characterize the liver transcriptome and apply single-cell RNA sequencing (scRNA-seq) for germline-depleted gonads. We reveal a transcriptional response that is distinct to germline depletion, including an overlap with mammalian pro-longevity interventions. Accordingly, we find that germline depletion can significantly promote longevity in a male-specific manner, increasing maximal lifespan by ∼50%.

To explore evolutionary conservation, we demonstrate that fine tuning of *eef1a1* (eukaryotic translation elongation factor 1 alpha 1), a differentially expressed candidate gene in germline-depleted killifish, is required for the longevity of germline-depleted *C. elegans*. Finally, we use multi-omics and physiological assays to demonstrate that old germline-depleted males experience prolonged healthspan through enhanced metabolic functions. Taken together, the observed sexual dimorphism supports the notion that classical evolutionary tradeoffs could be uncoupled and involve a possible systemic effect. Our findings propose that distinct flavors of the reproductive state can have a profound impact on vertebrate life-history.

### Reconstruction of the killifish gonad by scRNA-seq

As a relatively new vertebrate model, the cell types that constitute the killifish reproductive system and their corresponding molecular markers are largely unknown. Furthermore, determining gonadal composition is imperative for downstream analysis such as how distinct gonadal cell types respond to perturbations of the reproductive state. We therefore collected single-cell transcriptomes from one-month-old testis and ovaries to examine the gonadal composition and cell types involved. This age was chosen because killifish have already gone through sexual maturation at this time, and all cell types should be present. Gonads from three male and three female fish were dissociated enzymatically, and single-cell droplet barcoding was performed using the InDrop approach^44,45^ (**Figure 1a**). Following quality control, we recovered a total of 5695 single-cell transcriptomes. As the first step, the cells were clustered using uniform manifold approximation and projection (UMAP) and the Leiden algorithm, and batch correction was performed by applying the Harmony tool^46^ (see **Methods**). Finally, a distinction between the germline (gray) and somatic cells (blue) was made possible by using conserved germline markers (**Figure 1b**, top left, **Extended Data Figure 1a**, and see **Methods**).

**Figure 1:**
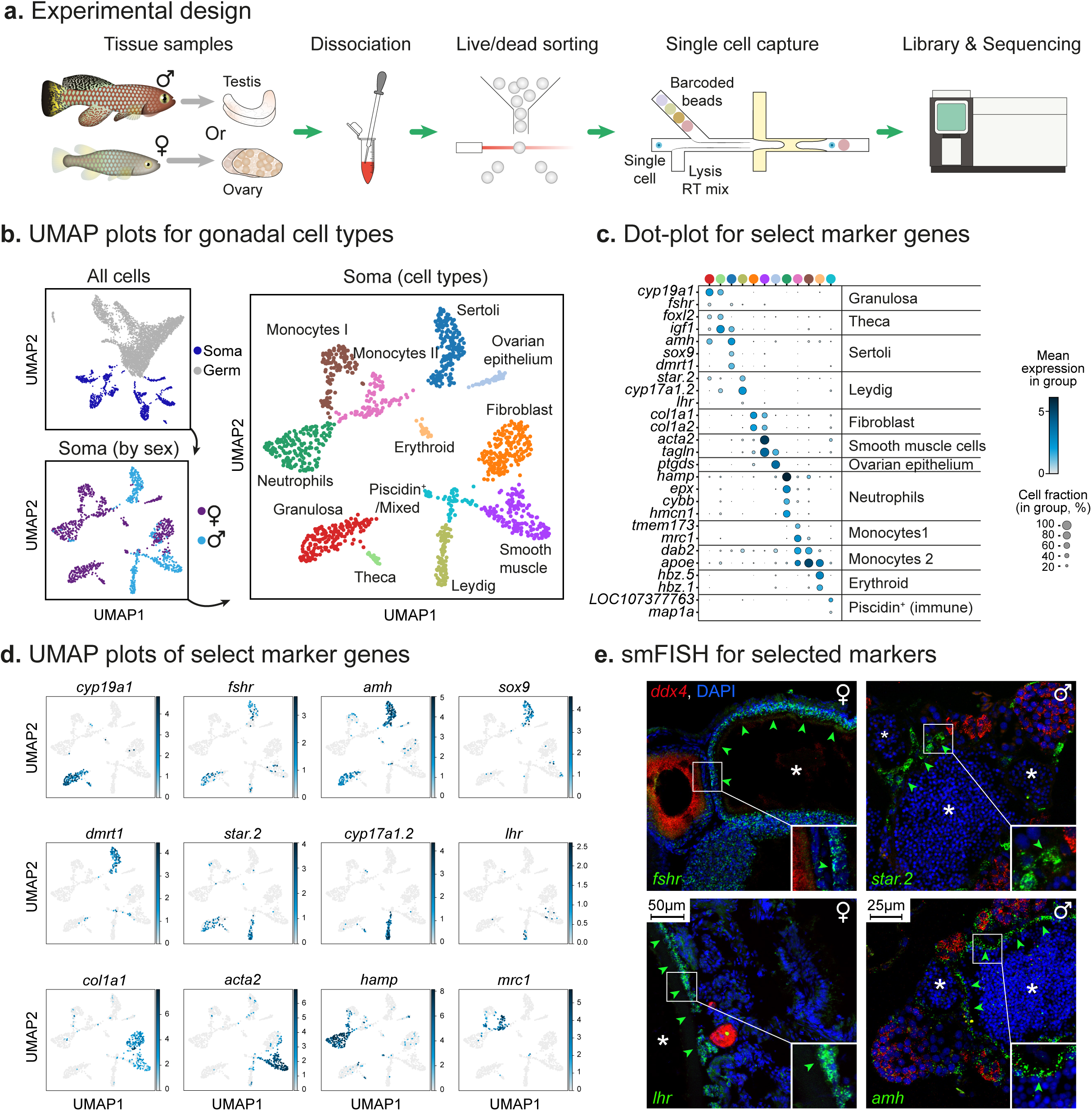
Reconstruction of somatic cell types in the gonads of the turquoise killifish. (**a**) Experimental workflow for characterizing male and female killifish gonadal cell types using InDrop single-cell RNA sequencing. (**b**) UMAP analysis of 5695 single cells from the testis and ovary of one-month-old fish. Clusters are color-coded either by somatic- and germ-cells (top left), by somatic cells from males and females (bottom left), or by somatic cell types (right). (**c**) Dot-plot representing the relative expression of select cell-type marker genes in each cluster. Clusters are color-coded according to (**b**, right). Some markers are shared between cell types, such as cells with similar functions (e.g. *amh* in the supporting Sertoli and Granulosa cells). A full list of cell-type markers can be found in **Supplementary Table S2**. (**d**) Gene expression UMAP plots of selected marker genes that correlate with primary cell types. (**e**) smFISH in ovaries (left) and testes (right) for selected markers. Marker for immature germ-cells (*ddx4*) in red (dot-plot in Figure 2d), and for somatic cell types *(fshr, star, lhr,* and *amh)* in green. Asterix highlights mature germ cells, which are *ddx4* negative. Green arrowheads highlight expression patterns. Representative of n ≥ 6 individuals. Scale bar: 50 µm for ovary, and 25 µm for testis. Note that individual channel images are available in source data for accessibility to colorblind readers.

### Distinct cell populations in male and female gonads

Vertebrate ovaries and testes are composed of several primary cell types, including supporting, steroidogenic, interstitial, and germ cells. In females, the supporting Granulosa cells are in direct contact with the oocytes, with a layer of steroidogenic Theca cells on top. In males, the supporting Sertoli cells enclose the germ cells, while the steroidogenic Leydig cells are located in clusters spread across the interstitium. To focus on the somatic gonad, we first filtered out the germline (**Figure 1b**, bottom left), grouped the somatic cells into 12 clusters (**Figure 1b**, right), and characterized their cell types based on the expression of conserved markers^42,43,47–56^ (**Figure 1c, d**). A list of all markers associated with each cluster and a list of specific killifish paralogues, as well as a list of markers which are conserved with zebrafish are available in **Supplementary Tables S1, S2**.

We then investigated the spatial architecture of the killifish gonad. Specifically, we selected evolutionary conserved cell-type-specific markers for the somatic compartment^43,49–52^ (**Figures 1c, d**), and performed single molecule fluorescent in-situ hybridization chain reaction (smFISH HCR, **Figure 1e**). These markers included *star* (steroidogenic acute regulatory protein, for Leydig cells), *amh* (anti-mullerian hormone, for Sertoli cells), and the lowly expressed gonadotropin receptor *fshr* (follicle stimulating hormone receptor for Granulosa cells). Additionally, we included a classical germ cell marker^57^ as a reference (*ddx4/vasa*, dead-box helicase 4).

Similarly, to characterize the germline compartment, we performed a sub-clustering analysis for the germ cell populations, which included 4294 male sperm cells, and only 74 female oocytes (**Extended Data Figure 1b**, left). The difference in capture efficiency can be attributed to the larger size and relative scarcity of female oocytes, which tends to exclude them from our droplet-based pipeline. Therefore, we focused primarily on the male germline, removing female-derived cells from our analysis, as well as a small cluster identified as erythroid cells (**Extended Data Figure 1b**, right, and see **Supplementary Table S3**).

Similar to the somatic cells, predictions of cell-type specific markers for the germline were made based on evolutionary conservation^43,49–52^ (**Figure 2a**) and a pseudotime analysis (see **Methods**, and **Extended Data Figure 2a**). To refine specific developmental stages, we further performed a comparison with similar single-cell data from the zebrafish male germline^43^ (**Figure 2b**), and describe predicted stage-specific markers (**Figure 2c-d**). One remaining cluster seemed to be of a mixed unspecified germline population (**Figure 2a-d**). Specifically, using these analyses we identify primary spermatogenesis stages, encompassing spermatogonia (SPG), spermatocytes, and spermatids, along with several sub-classifications. For example, both spermatogonial clusters (2 and 5), were correlated with undifferentiated and differentiated spermatogonia in a parallel zebrafish dataset^43^, SPG and SPG2, respectively (**Figure 2a-d**). The full list of differentially expressed genes of each germline cluster is presented in **Supplementary Tables S1, S3**.

**Figure 2:**
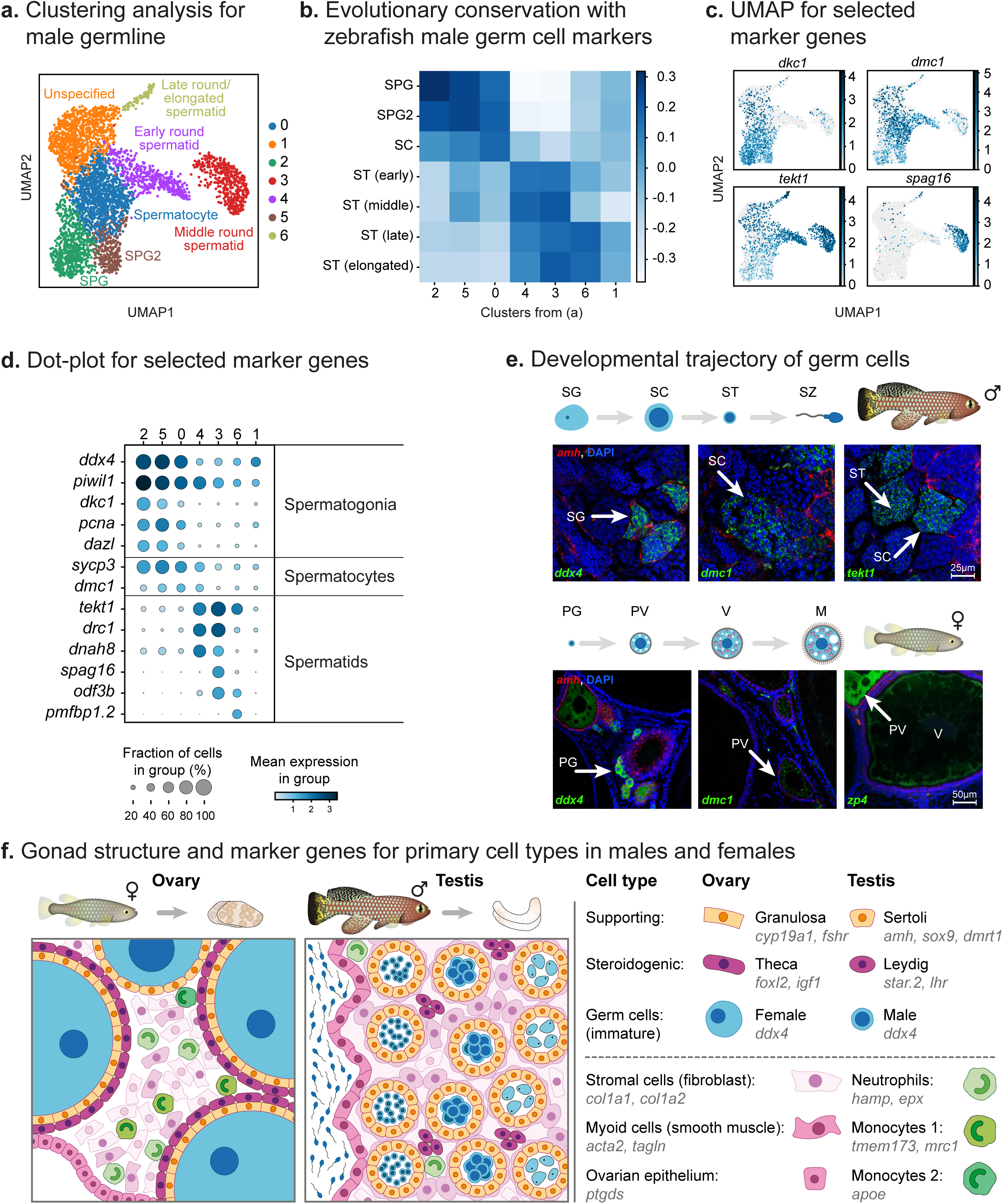
Reconstruction of germ cell types in the gonads of the turquoise killifish. (**a**) Sub-clustering of the male germ cells excluding the female germ cells and erythroid cells. Cells are color-coded and labeled by cell type. (**b**) Heatmap demonstrating the correlation between differentially expressed genes for each germline cluster from (**a**, X-axis), compared to distinct cell types identified in the zebrafish male germline (Qian et al^43^, Y-axis). (**c**) Selected gene expression UMAP plots of male germ cell marker genes. A full list of all markers can be found in **Supplementary Table S3**. (**d**) Dot-plot representing the relative expression of select cell-type specific marker genes in each cluster. Clusters are numbered according to (**a**). (**e**) A model of germ cell development in killifish (top), in either males (top) or females (bottom) (modified from^28^). Bottom: smFISH for selected markers in testis (top) or ovary (bottom), including markers for supporting cells (*amh*, in red), and for stage-dependent germ cell markers (*ddx4, dmc1, tekt1*, and *zp4*, in green). Representative of n ≥ 6 individuals. Scale bar: 25 µm (top) and 50 µm (bottom). SG: spermatogonia; SC: spermatocytes; ST: spermatids; SZ: spermatozoa; PG: primary growth; PV: pre-vitellogenic; V: vitellogenic; M: Mature. (**f**) A predicted model illustrating the spatial distribution of primary gonadal cell types in killifish ovary and testis, and their associated marker genes. Shapes of sperm cells indicate different developmental stages, from early stages (right) to fully differentiated sperm (left). See detailed differentiation trajectory in (**e**, top).

We next visualized selected markers of male germ cells using smFISH, including *ddx4* (spermatogonia), *dmc1* (DNA meiotic recombinase 1, spermatocytes), and *tekt1 (*tektin-1, primarily in spermatids, **Figure 2e**, top). Even though female germ cells were scarce in our data, we were able to predict several markers genes according to the association of female cells with male clusters (**Extended Data Figure 1b**). We further validate several prediction using smFISH, including *ddx4*, *dmc1*, and *zp4* (zona pellucida 4) (**Figure 2e**, bottom). Combining single-cell transcriptomics with smFISH made it possible to identify somatic- and germline-derived gonadal cell types in killifish.

Overlaying the identified marker genes with teleost gonadal architecture^58–60^ allows us to propose a model for the male and female killifish gonads (**Figure 2f**). These data provide the foundations for exploring the killifish reproductive state. For example, distinct molecular markers are important to better evaluate cell-type-specific responses to perturbation via a transcriptional signature of a distinct cell type, or changes in cell composition.

### Perturbation of the HPG reproductive axis in killifish

The hypothalamic-pituitary-gonadal axis (HPG) plays a central role in controlling vertebrate puberty. Specifically, the follicle stimulating hormone (FSH) and luteinizing hormone (LH) are released by the pituitary, bind to their gonadal receptors, and support ovary and testis development. In turn, steroid hormones are secreted from the gonads, and feed back to the brain to fine tune hormone levels (**Figure 3a**). Although the HPG is largely vertebrate-conserved, there are subtle differences between fish and mammals, which give rise to altered regulatory networks^61^.

**Figure 3:**
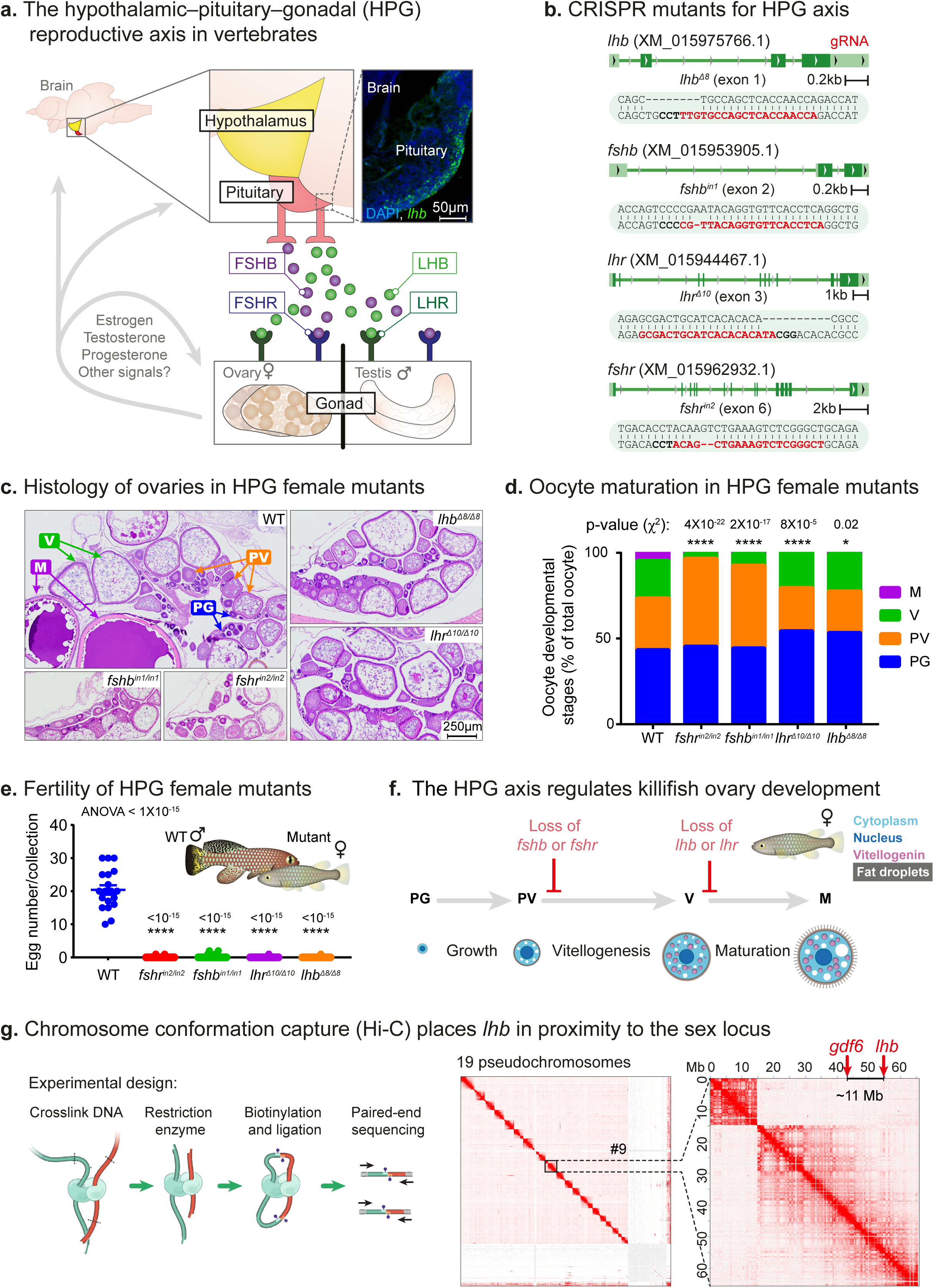
Genetic perturbation of the reproductive axis in killifish. (**a**) Schematic illustration of the vertebrate hypothalamic-pituitary-gonadal axis (HPG): FSH and LH are released from the pituitary, and travel through the bloodstream to their target receptors in the gonad to stimulate germ cells production. Top right: an immunostaining for *lhb* (green) and DNA (blue) in histological section of the pituitary in the turquoise killifish. Representative of n =3 individuals. Scale bar: 50 µm. (**b**) Generation of HPG CRISPR mutants, depicting the gRNA targets (red), protospacer adjacent motif (PAM, in bold), and indels. (**c**) Representative histological sections, depicting ovaries from one-month-old females of the indicated genotypes. n ≥ 5 individuals from each genotype. Scale bar: 250 µm. Oocyte development stages: PG: primary growth; PV: pre-vitellogenic; V: vitellogenic; M: Mature. (**d**) Quantification of oocyte maturation in each of the genotypes from (**c**). Data presented as proportion of each oocyte developmental stage. n ≥ 5 individuals for each experimental group. Significance was measured by a two-sided χ^2^ test with the WT proportion as the expected model and FDR correction. P-values are indicated. Abbreviations represent the oocyte developmental stage according to^102^. (**e**) Quantification of female fertility output. Each dot represents the number of eggs per breeding pair of the indicated genotypes, per week. A total of 6 independent pairs and 4 independent collections were performed. Error bars show mean□±□SEM with individual points. Significance was calculated using one-way ANOVA with a Dunnet post-hoc compared to the WT and p-values are indicated. (**f**) A model for the role of gonadotropin signaling in controlling germ cell development in killifish females (modified from^28^). FSH controls follicle growth and the transition from pre-vitellogenic (PV) to vitellogeinc (V), while LH is important for oocyte maturation (M). The specific mutation inhibits the relevant processes. (**g**) Experimental workflow for Hi-C (left). A genome-wide contact matrix (center), demonstrating the 19 linkage groups (pseudochromosomes) that correspond to the 19 killifish chromosomes. The submatrix (right) corresponds to interactions on pseudochromosome 9 (chromosome 5). Locations of the *lhb* gene and the sex determining locus (growth differentiation factor 6a, *gdf6a*) are indicated (red arrows).

As the first step in manipulating killifish reproduction, we genetically perturbed the HPG reproductive axis by mutating genes that correspond to primary pituitary hormones (FSH and LH), and their corresponding gonadal receptors (FSHR and LHR, respectively). Specifically, using CRISPR/Cas9 genome editing, followed by outcrossing for several generations to reduce off-target effects, we produced the following homozygous mutants: *fshb^in^*^1^*^/in^*^1^, *fshr^in^*^2^*^/in^*^2^*, lhb*^Δ8^*^/^*^Δ8^, and *lhr*^Δ10^*^/^*^Δ10^ (**Figure 3b**). Germline maturation was assessed by staining tissue sections with Hematoxylin and Eosin (H&E), and scoring the proportion of specific developmental stages^59,60^, in both females (**Figures 3c, d**) and males (**Extended Data Figure 3a, b**). Our findings demonstrate that, as in zebrafish and medaka^62^, female germ cell maturation is arrested at distinct stages in response to mutating *fshb, lhb*, or their corresponding receptors, and these mutants are therefore largely infertile (**Figure 3e**).

In fish, one of the main energy and metabolic investments during female reproduction is in vitellogenesis^63^; the resource-intensive process of yolk formation via nutrient deposition into the oocyte. Our results indicate the *fshb^in^*^1^*^/in^*^1^ and *fshr^in^*^2^*^/in^*^2^ female mutants display a striking increase in the proportion of previtellogenic eggs (stage PV), at the expense of vitellogenic and mature eggs (stages V and M, respectively, **Figure 3d**). No effect was observed in oocytes during the early developmental stages of primary growth (stage PG). This observation implies that oocyte maturation is arrested before vitellogenesis. Conversely, female mutants for *lhb* and its receptor *lhr* exhibit an increase in vitellogenic oocytes (stage V), but a decrease in mature oocytes (stage M). This, suggests that oogenesis is arrested in these mutants at a later stage, before final maturation (**Figure 3d**). There is no significant effect on male growth or germline development, and all mutants (aside from the male *lhb*^Δ8^*^/^*^Δ8^, which is discussed below) were produced at the expected Mendelian ratios (**Extended Data** Figures 3a-e).

Together, our results reveal the developmental windows in which gonadotropins are required for specific stages of female killifish germline differentiation (summarized in **Figure 3f**). However, even though several theories predict that perturbing reproduction could have a tradeoff with other life-history traits, such as growth, none of the HPG mutants exhibited any significant effect on somatic growth (**Extended Data Figure 3d**).

### Hormonal loss-of-function can be phenotypically rescued

All HPG knockout females are largely infertile (**Figure 3e**). To test whether the phenotypes in *lhb*^Δ8^*^/^*^Δ8^ mutants are directly mediated by the loss of hormone function, we attempted to restore fertility experimentally. Specifically, we used a platform for functional interrogation of peptide hormones, which we have recently developed for killifish^38^, to ectopically express hormones in muscle tissue (**Extended Data Figure 3f**, left). These hormones are then predicted to be secreted into the circulation and ultimately reach the gonads. *Lhb* was cloned from turquoise killifish cDNA and tagged with GFP separated by the T2A self-cleaving peptide, which leaves the hormone untagged after cleavage.

This approach allowed us to restore oocyte differentiation and female fertility to *lhb*^Δ8^*^/^*^Δ8^ mutants (**Extended Data Figure 3f**, right). A similar rescue was also described in *fshb^in^*^1^*^/in^*^1^ as part of the method development^38^. These results suggest that the observed reproductive phenotypes are indeed directly attributed to a loss of hormone function. Our conclusions are further supported by the undetected transcript levels of the receptor in the corresponding *lhr* and *fshr* homozygous mutants, as evaluated by smFISH (probably due to nonsense-mediated mRNA decay, **Extended Data Figure 3g**).

### Chromosome conformation capture places *lhb* near the sex locus

Curiously, although most HPG mutants were produced at the expected Mendelian ratios, *lhb*^Δ8^*^/^*^Δ8^ mutant males were rarely observed (**Extended Data Figure 3c**). This raised the possibility that the gene is located near the killifish sex determination gene *gdf6*^34^ (growth differentiation factor 6) on chromosome 5, and therefore, is not equally segregated between males and females. Since *lhb* (LOC107396183) is currently on an unplaced genomic scaffold, we performed chromosome conformation capture (Hi-C^64^) and used it to scaffold the killifish genome. This indeed demonstrated that the *lhb* gene is located only 11 Mb away from the sex locus (**Figure 3g** and see **Methods** for details). The Hi-C raw data and analysis are available in the GEO database (see **Methods**). Additionally, by generating 814,717,494 Hi-C reads, our analysis also corrects many large-scale errors in the killifish genome and assists in positioning short contigs within chromosomes (improving the N90 from 152,787 to 298,000 and reducing scaffold numbers from 5,897 to 2,614, **Extended Data Figure 3h**).

### Germline depletion enhances female somatic repair

One prediction of the ‘disposable soma theory’ is that perturbing reproduction should have a tradeoff with other physiological traits, particularly somatic damage repair^5,6^. To test this, we selected the infertile *fshr^in^*^2^*^/in^*^2^ and *lhr^Δ^*^10^*^/Δ^*^10^ mutants, in which oocyte development is blocked at early stages, either before the resource-intensive vitellogenesis process or before oocyte maturation, respectively (**Figure 3c-d**). The alternative ‘endocrine signals’ hypothesis suggests that physiological traits will be altered only by completely removing the germline (while the somatic gonad remains intact). Therefore, to simulate this physiological scenario we mutated the *dnd1* gene (dead end^65^) to produce a germline-depleted killifish model (**Figure 4a**, left).

**Figure 4:**
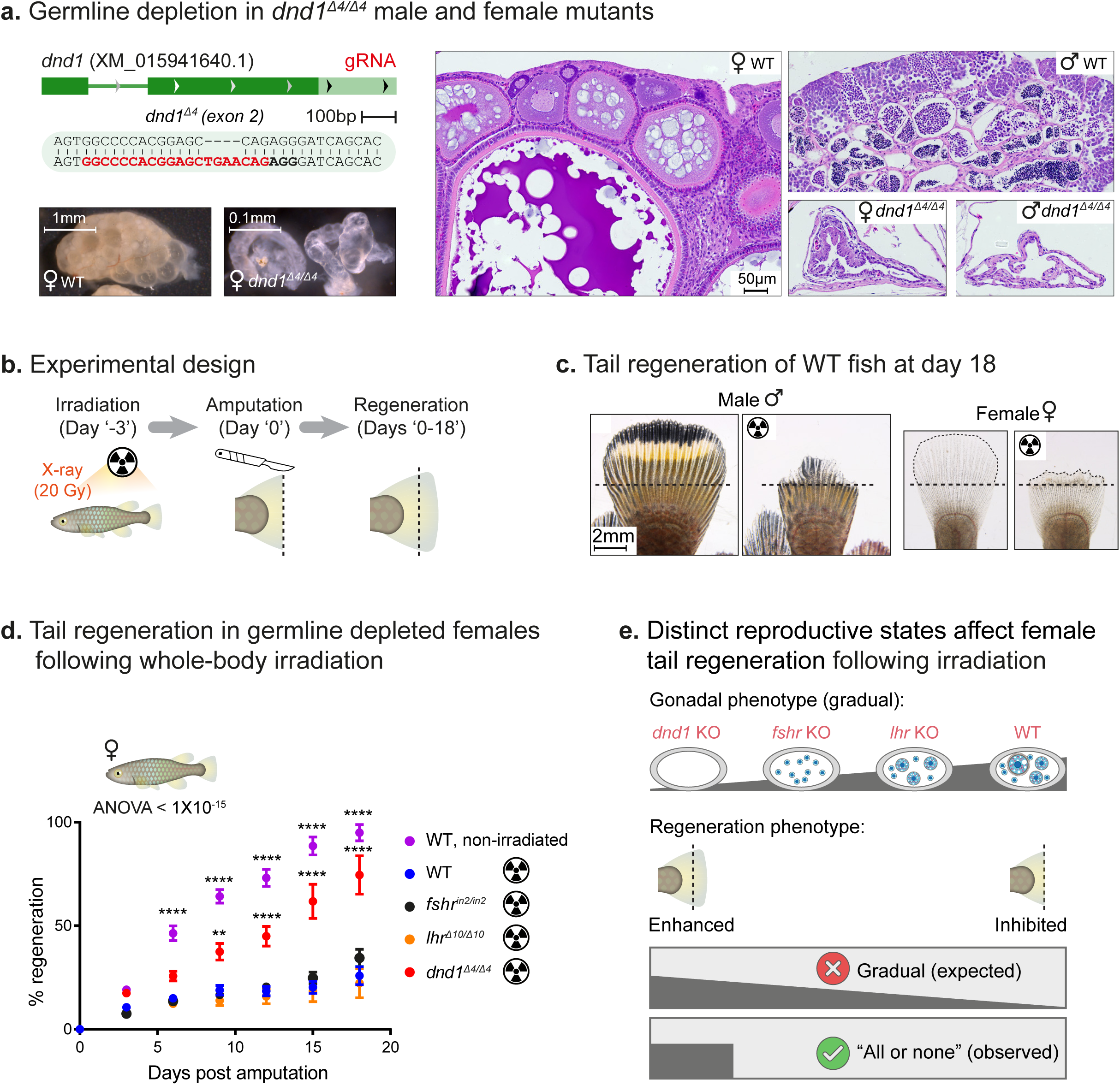
Germline depletion enhances somatic regeneration in females. (**a**) Top left: generation of a CRISPR mutant for the *dnd1* gene, depicting the gRNA target (red), PAM (in bold), and indel. Bottom left: images of female gonads from one-month-old fish, either WT or *dnd1*^Δ4^*^/^*^Δ4^ mutants. Scale bar: 1 mm for WT fish, 0.1 mm for *dnd1*^Δ4^*^/^*^Δ4^ mutant fish. Right: histological sections of male and female gonads from one-month-old fish, either WT or *dnd1*^Δ4^*^/^*^Δ4^ mutants. Representative of n ≥ 4 individuals from each genotype. Scale bar: 50 µm. (**b**) Experimental workflow for assessing tail-fin regeneration following irradiation. Tail fish amputation was performed three days after irradiation, and regrowth was measured every three days for a total of 18 days. (**c**) Examples of regenerated tails at day 18 post amputation, in male (left pair) or female (right pair), under either control (left of each pair) or irradiated (right of each pair) conditions. The black dashed line represents the amputation plane. Scale bar: 2 mm. (**d**) Tail-fin regeneration following whole-body irradiation in females. Genotypes and treatments are indicated (n =14 individuals for non-irradiated controls, 13 for irradiated WT, *lhr*, and *dnd1*, and 12 for *fshr* fish). The length of the outgrowth was calculated as the percentage of the original fin size (before the amputation). Error bars indicate mean□±□SEM. Significance was calculated using repeated-measures two-way ANOVA with a Dunnet post-hoc compared to the irradiated WT, and p-values are indicated. (**e**) A model summarizing the effect of our genetic perturbations on regenerative potential following irradiation. While a gradual effect is observed on germline differentiation, an “all or none” effect is seen in regeneration potential.

*Dnd1* was previously shown to be essential for germline formation in fish, because it is responsible for regulating primordial germ cell migration into the embryonic somatic gonad^65^. Accordingly, *dnd1*^Δ4^*^/^*^Δ4^ homozygous killifish mutants are germline-free (**Figure 4a**, right). Similar to the HPG mutants described above, *dnd1*^Δ4^*^/^*^Δ4^ fish displayed no significant tradeoff with somatic growth (**Extended Data Figure 4a, b**). Germline depletion in either zebrafish or medaka causes an all-male sex-reversal^39,40^, which precludes the study of sexual dimorphism (particularly of females). However, both male and female germline-free *dnd1*^Δ4^*^/^*^Δ4^ killifish were obtained at the overall expected Mendelian ratios (**Extended Data Figure 4c**). Thus, demonstrating that much like in mammals, killifish undergo germ-cell-independent sex differentiation.

To evaluate the effect of *dnd1*^Δ4^*^/^*^Δ4^*, fshr^in^*^2^*^/in^*^2^, and *lhr*^Δ10^*^/^*^Δ10^ on somatic damage repair, we assessed tail regeneration capacity following irradiation as a proxy for a physiological challenge^7^. Killifish were first exposed to a sub-lethal level of whole-body ionizing radiation (X-ray, 20 Gy), and regenerative potential was assessed by measuring the rate of tail-fin regrowth following amputation (**Figure 4b, c**). Interestingly, while the homozygous *lhr*^Δ10^*^/^*^Δ10^ and *fshr^in^*^2^*^/in^*^2^ mutations had no apparent effect, regeneration was significantly enhanced in germline-free *dnd1*^Δ4^*^/^*^Δ4^ fish, specifically in females (**Figure 4d, Extended Data Figure 4d**). We did not test HPG mutants in males (**Extended Data Figure 4d**), as they had no germline phenotype (**Extended Data Figure 3a, b**).

The *dnd1*^Δ4^*^/^*^Δ4^ mutation did not enhance tail regeneration in old fish (**Extended Data Figure 4e**), suggesting that beneficial effects might only be obtainable in response to a particular challenge. Our results indicate that only specific reproductive defects, namely germline depletion, can enhance somatic maintenance in a sex-specific manner (an ‘all or none’ effect, see model in **Figure 4e**). This suggests that the observed phenotypes may not be limited to a classical tradeoff, which predicts a more gradual phenotype that is inversely correlate with the resources invested in the germline (**Figure 4e**).

Recent findings have suggested that once the germline is damaged by irradiation, its ‘expensive’ repair can have a direct tradeoff with somatic regeneration^7^. Thus, offering an interesting prediction, proposing that irradiating the entire body (both the tail and the germline) would have a different outcome when compared to tail-specific irradiation. To compare these two possibilities, we exposed WT fish to the same dose of radiation, either whole-body or tail-specific irradiation to avoid damaging the gonad (**Figure 5a** and **Extended Data Figure 5a**). Even though only whole-body irradiation induced extensive damage to the germline, both interventions equally inhibited tail regeneration (**Figure 5b**). Thus, our data suggest that the resources required to repair the damaged germline might not be the primary source for the observed phenotypes, but rather the direct damage the tail has sustained.

**Figure 5:**
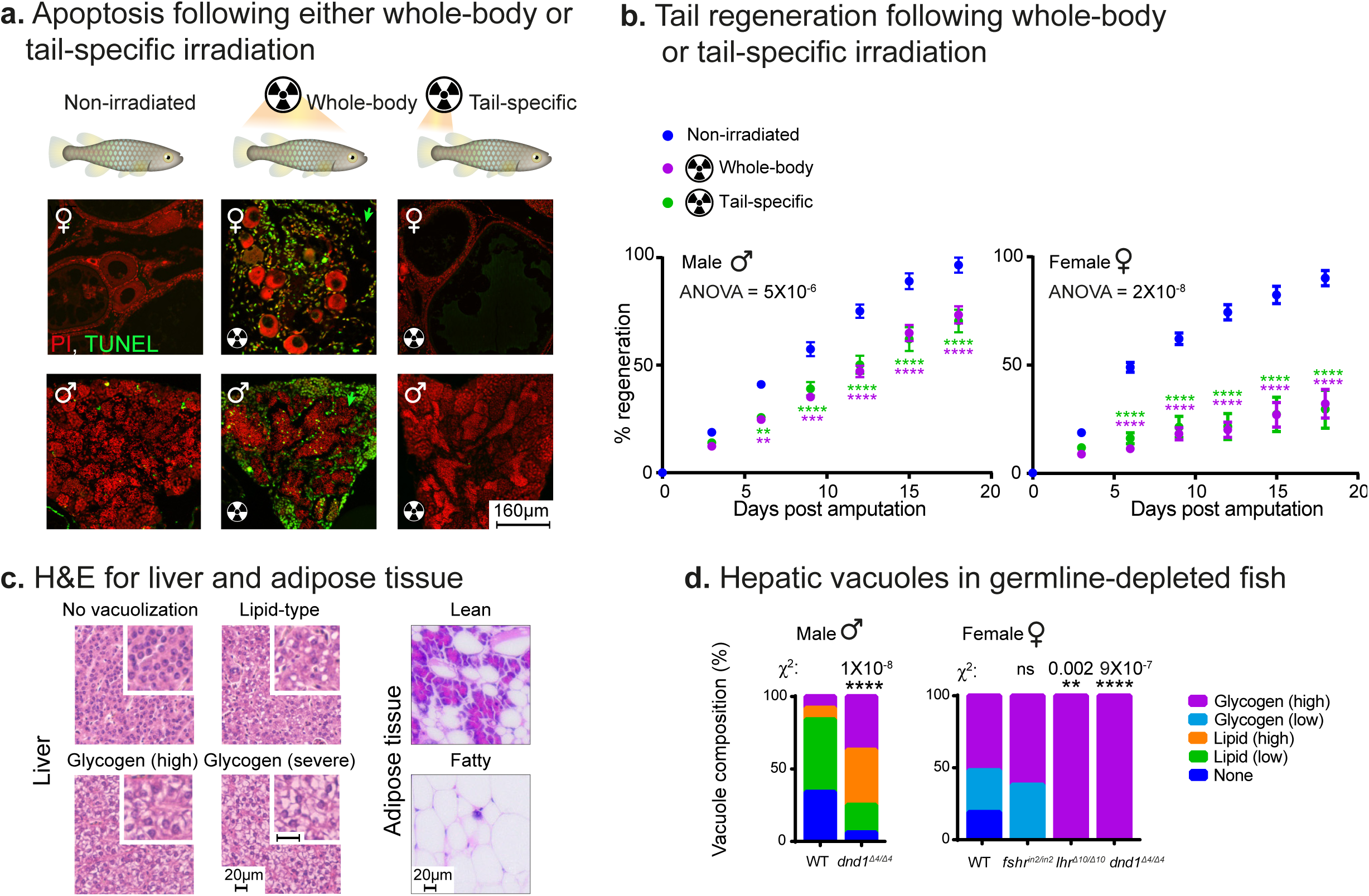
Exploring tradeoffs between reproduction and DNA damage repair. (**a**) Representative images of apoptosis detected by the TUNEL assay (green and green arrows) and DNA (PI, in red). Assay was performed in tissue sections of ovaries (top) or testis (bottom), under either control conditions (left), following whole-body irradiation (center) or tail-specific irradiation (right). n≥ 6 individuals in each experimental group. Scale bar: 160 µm. (**b**) Tail-fin regeneration following whole-body or tail-specific irradiation in males (left) and females (right). Treatments are indicated (n = 11 individuals for non-irradiated females, 18 for whole body irradiated females, 19 for tail-specific irradiated females, 20 for non-irradiated males, 21 for whole body irradiated males and 29 for tail-specific irradiated males). Error bars indicate mean□± SEM. Significance was calculated using repeated-measures two-way ANOVA with a Tukey post-hoc, and p-values are indicated. The significance of each experimental group is color-coded and relates to the comparison to the non-irradiated fish. (**c**) Representative histological sections depicting livers (left) and adipose tissue (right) from one-month-old fish with the indicated vacuole composition or adiposity. Scale bar: 20 µm (**d**) Quantification of liver vacuole composition, according to (**c**), in male (left) and female (right) fish of the indicated genotypes. Data presented as proportion of each vacuole-type. 3 histological sections from n ≥ 5 individuals for each experimental group. Significance was measured by two-sided χ^2^ test with FDR correction, with the WT proportion as the expected model. P-values are indicated.

Finally, to explored other physiological changes we examined the liver, a primary metabolic organ that is involved in the energy production required for vitellogenesis. Specifically, we characterized the accumulation of hepatocellular vacuolizations^66,67^, which are either lipid-type (round, sharp edges) or glycogen-type (irregular, lightly stained) (**Figure 5c**). Our results indicated no strong correlation between the severity of the gonadal phenotype and the metabolic effect in the liver (**Figure 5d**) or in the internal visceral adipose tissues^68^ (**Supplementary Table S4**). Therefore, to better understand how germline depletion affects the surrounding somatic cells we turned to scRNA-seq.

### scRNA-seq of *dnd1*^Δ4^*^/^*^Δ4^ gonads suggests sex-specific effects

To explore differential gene expression between WT and germline free fish we collected single-cell transcriptomes from one-month-old testis and ovaries of *dnd1*^Δ4^*^/^*^Δ4^ fish (**Figure 6a, b**). Germline-depleted gonads are naturally smaller compared to WT (**Figure 4a**, as also seen worms^8^). This required samples to be pooled to provide enough cells (see **Methods**). Following quality control, we recovered an additional 4361 single-cell transcriptomes from the gonads of *dnd1*^Δ4^*^/^*^Δ4^ mutants (2539 from males, and 1822 from females). Male and female samples were analyzed separately, and UMAP and batch correction were applied as described above. Cells were then grouped into 11 and 18 clusters, in males and females, respectively (**Figure 6a, b** and **Supplementary Tables S1, S5, S6**).

**Figure 6:**
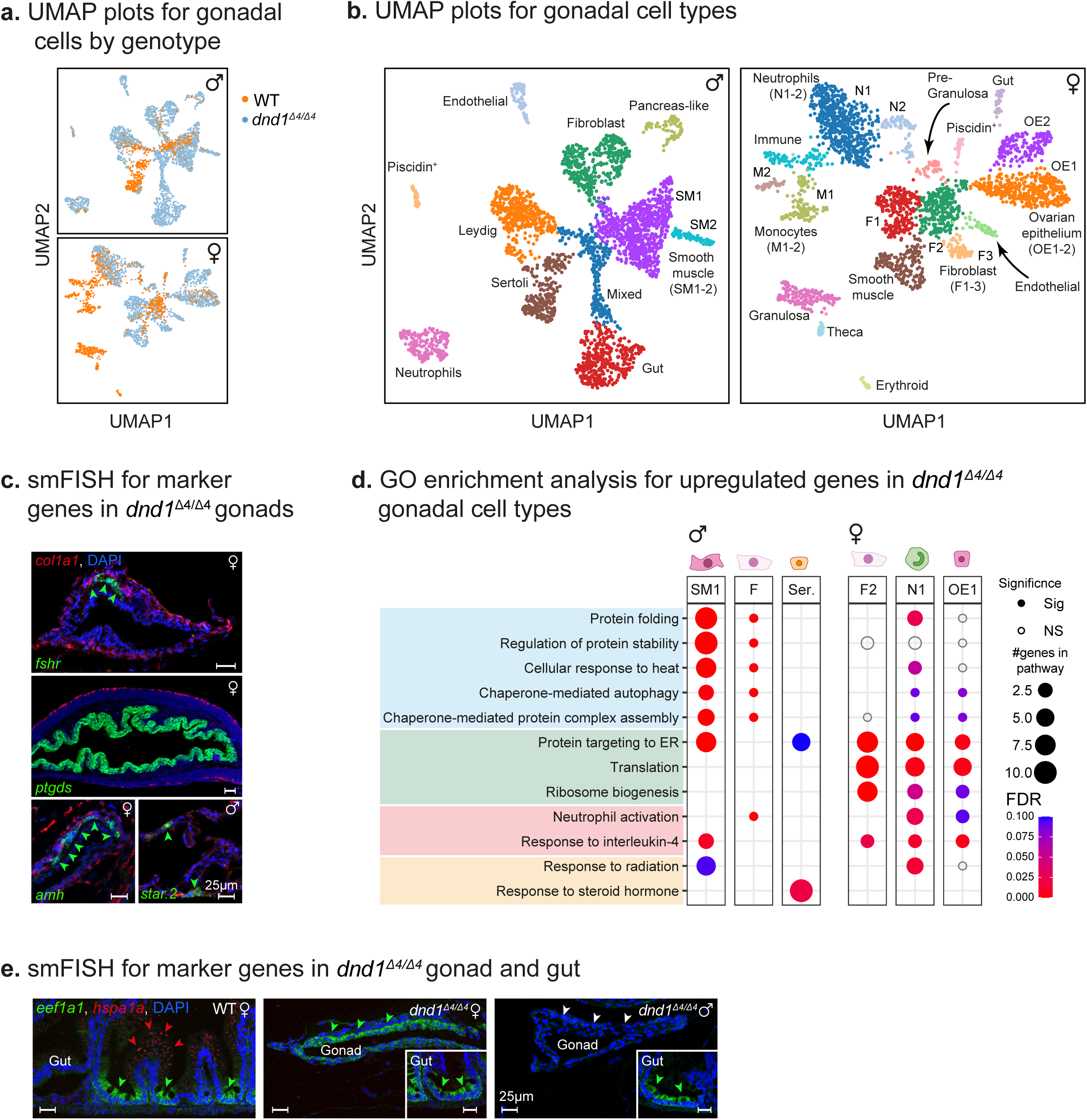
Single-cell RNA sequencing reveals a sex-specific somatic response to germline depletion. (**a**) UMAP of cells from WT and *dnd1*^Δ4^*^/^*^Δ4^ mutant gonads (color-coded by genotype), separated into males (top) and females (bottom). (**b**) UMAP from (**a**), color-coded by cell types. Cell-type markers are listed in **Supplementary Tables S1, S5, S6**. (**c**) smFISH in ovaries and testes (as indicated by sex symbols) for selected markers in *dnd1*^Δ4^*^/^*^Δ4^ mutants. Markers for fibroblasts (*col1a1*) in red, and for other somatic cell type markers (*fshr, star, amh,* and *ptgds*) in green (as described in Figure 1e). Green arrowheads highlight expression patterns. Representative of n ≥ 6 individuals. Scale bar: 25 µm. (**d**) Enrichment analysis (GO) for upregulated genes in *dnd1*^Δ4^*^/^*^Δ4^ mutants at the indicated clusters/cell types. Distinct patterns between males and females are observed. GO enrichment was called at FDR < 5%. (**e**) smFISH for selected markers in gut, including the intestinal crypts and villi as controls (top, and small insets), ovaries (center), and testes (bottom) in *dnd1*^Δ4^*^/^*^Δ4^ mutants. A marker for translation initiation (*eef1a1*) in green, and for heat shock response (*hspa1a1*) in red. Green and red arrowheads highlight expression patterns. White arrowhead indicates lack of expression. Small insets highlight the intestinal crypts, as controls. Representative of n ≥ 10 individuals. Scale bar: 25 µm.

While germline lineages were completely missing in the homozygous *dnd1*^Δ4^*^/^*^Δ4^ gonads, primary somatic cell types could be assigned and visualized using smFISH (**Figures 1c-d, 6a-c, Extended Data Figure 6a, b**). Some interesting differences in cell composition were observed, such as a significant shift in immune cells in the female gonads. Specifically, we observed that while both monocytes and neutrophils are present in the WT gonads (19.3% and 15.7% of total cells, respectively), in *dnd1*^Δ4^*^/^*^Δ4^ mutants we could only detect neutrophils (28.6% of total cells, with no monocytes present. p < 0.0001, Fisher’s exact test for comparing the changes in cell-type numbers, **Figure 6a, b**). Other somatic cell types, such as fully differentiated Granulosa and Theca cells were not detected in the *dnd1*^Δ4^*^/^*^Δ4^ gonads (**Figure 6a, b**). This is either due to relatively small numbers of these particular cell types or because the terminal differentiation of pre-Granulosa to Granulosa and Theca was shown to require interaction with oocytes^69^. We confirmed these findings using smFISH in *dnd1*^Δ4^*^/^*^Δ4^ mutant females, demonstrating that while we were able to detect *amh* (a pre- and mature Granulosa marker), we failed to detect *cyp19a1* in the gonad (aromatase, a marker for mature Granulosa, **Figures 6c, Extended Data Figure 6c**).

To identify molecular pathways that are enriched following germline depletion, we focused on selected clusters that contained cells from both genotypes (**Figure 6a, b**), including fibroblasts (in both sexes), neutrophils and ovarian epithelium (in females), and Sertoli and smooth muscle cells (in males). We then screened each cell type for genes that are differentially expressed between WT and *dnd1*^Δ4^*^/^*^Δ4^ mutant cells and conducted pathway enrichments using gene ontology (GO, conducted separately for males and females, **Figure 6d**). We decided to focus on upregulated genes, which correspond to most of the differentially expressed genes in the mutant cells (**Supplementary Table S7**).

Our results highlighted a striking sexual dimorphism in the response to germline depletion. For example, the upregulated genes in female mutants appeared to be associated with translation and ribosome biogenesis (**Figure 6d**). Candidate genes include *eef1a1* (**Extended Data Figure 6d**), which was upregulated in females, while in male mutants it was downregulated. Using smFISH, we further confirmed that *eef1a1* was highly expressed in *dnd1*^Δ4^*^/^*^Δ4^ ovaries compared to testis (using the intestinal crypts as controls, **Figure 6e**). A similar pattern was observed by focusing on ovarian epithelium, marked by *ptgds* (prostaglandin D2 synthase, **Extended Data Figure 6e**). These results are in agreement with the role of protein synthesis in regenerative processes, including neuronal regeneration^70^, stem cell function^71^, liver regeneration^72^, and DNA damage^73^. In contrast, the upregulated genes in male mutants appeared to be more related to stress resistance and protein homeostasis, which are classical pro-longevity pathways^74^.

### Comparing hepatic transcriptional signatures in fish and mice

Based on the single-cell transcriptional responses, we first set to explore whether a similar pro-longevity signature exists in other peripheral tissues. Therefore, we characterized the hepatic transcriptome of young fish (5 weeks old), either WT or germline-depleted. The differential expression analysis revealed a total of 2031 downregulated and 1665 upregulated genes (FDR of 0.05, **Supplementary Table S8**).

While many genetic, pharmacological, and dietary interventions can extend murine lifespan, we were curious to explore which classical longevity interventions in mice is most similar to germline depletion. Therefore, we compared our findings with a recent dataset that characterized the hepatic transcriptional responses to 8 different longevity interventions (in young male and female mice^75^, **Figure 7a**). Interventions included rapamycin, caloric restriction (CR, 40%), methionine restriction (MR, 0.12%), growth hormone receptor knockout (GHRKO), acarbose, 17-α-estradiol (17α-E2), Protandim, and Snell dwarf mice that are *Pit1* knockout (POU domain class 1 transcription factor 1).

**Figure 7:**
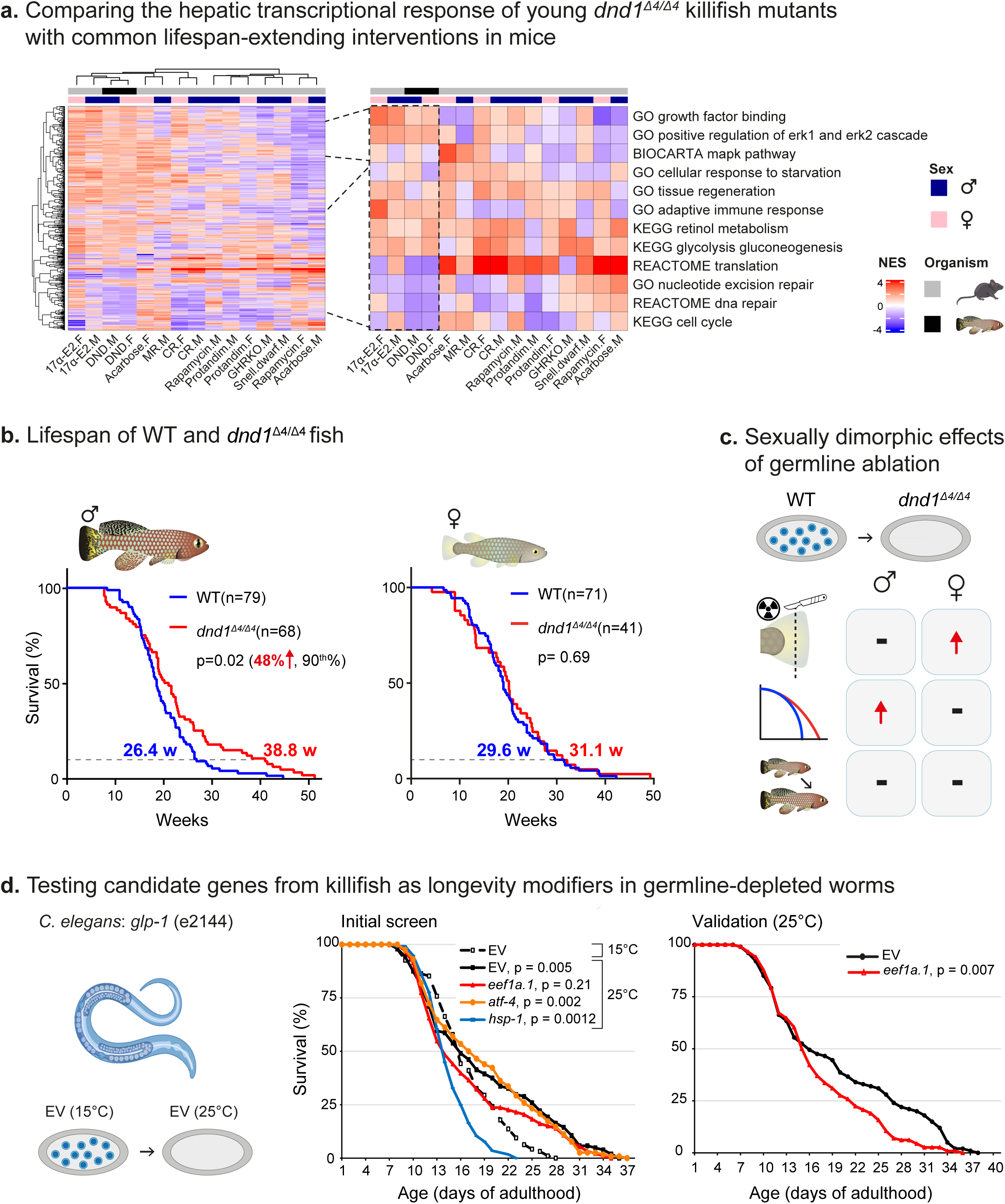
Germline depletion extends male lifespan. (**a**) Heatmap for normalized enrichment scores (NESs) in the liver, comparing the response to lifespan-extending interventions in mice with young *dnd1*^Δ4^*^/^*^Δ4^ mutant fish (left, selected pathways highlighted on the right) A full list of pathways can be found in **Supplementary Table S8**. (**b**) Lifespan of WT and *dnd1*^Δ4^*^/^*^Δ4^ fish, assessed separately for males (left) and females (right). P-values for differential survival in log-rank tests, maximal lifespan (measured at the 90^th^ percentile), and fish numbers are indicated. (**c**) A schematic model proposing that germline depletion (top) can regulate somatic regeneration (upper center) and longevity (lower center) in a sex-specific manner, with no significant tradeoff with somatic growth (bottom). (**d**) Lifespan of *glp-1*(e2144) mutants in *C. elegans*, grown either at the restrictive 25°C (germline depleted, sterile) or 15°C (permissive, fertile), and fed with the indicated RNAi. Experimental design is also presented (left). Initial mini-screen comparing all experimental conditions to baseline (fertile worms fed with an empty vector, EV, center). A validation experiment comparing EV and *eef1a1* RNAi fed sterile worms (right). P-values were calculated according to log-rank with an FDR correction, when required. Worm numbers and raw data can be found in **Supplementary Table S4**.

We then performed functional enrichment analysis for differentially expressed genes in germline-depleted young fish and compared them with enriched pathways in the mouse dataset^75^ (**Figure 7a** and **Supplementary Table S8**). Applying unsupervised clustering assisted us in identifying that germline depletion primarily clustered with 17α-E2, for both males and females (**Figure 7a**). Within this cluster, partial pathway overlaps include activation of ERK signaling (extracellular signal-regulated kinases), modified immune response, and carbohydrate metabolism. Other pathways were more enriched in one sex compared to the other, such as cellular response to starvation (male killifish), and tissue regeneration (female killifish). Our findings provide important insights into possible mechanisms of action that might be shared with mammalian longevity interventions.

### Germline depletion extends male lifespan

So far, the liver and single-cell data prompted further examination of the long-term effects of germline depletion on lifespan. To exclude other possible costs of reproduction, particularly mating and physical damage^76^, animals were single-housed. Excitingly, we observed a male-specific lifespan extension (**Figure 7b**, p = 0.02, log-rank test), while the lifespan of female *dnd1*^Δ4^*^/^*^Δ4^ mutants remained unchanged (p = 0.69, log-rank test). The longevity effect in males was not uniform across life, as the median lifespan increased by ∼11%, while the maximal lifespan by ∼50% (90^th^ percentile^77,78^).

We therefore investigated the shape of the lifespan curve by using a Quantile-Quantile (Q-Q) plot (**Extended Data Figure 7a**, left). In a hypothetical case where control and experimental groups are similar, the plot would be along a straight 45° angle line (as seen in females, **Extended Data Figure 7a**, right). This is true in our data only up to ∼80^th^ percentile (175 days). We validated this analysis by using Boschloo’s test^78^, identifying that while there is no significant lifespan difference at the 50^th^ and 75^th^ percentiles (p=0.223 and 0.12, respectively), a significant difference was observed at the 90^th^ percentile (p=0.04). These findings are consistent with the ‘neck-and-shoulder’ lifespan curve previously observed in germline-depleted worms^8^.

In agreement with our sex-dimorphic longevity phenotype (**Figure 7b**), 17α-E2 treatment in mice extends only male lifespan^79^. 17α-E2 is a naturally occurring enantiomer of 17β-estradiol (17β-E2), which appears to be non-feminizing due to minimal activation of canonical estrogen receptors^80^. Therefore, we next compared whether other common gonadal/hormonal interventions in male mice^75,81,82^ (e.g. castration, estrogen receptor KO, hormone replacement) might overlap with the transcriptional response of germline depletion. Our data indicated that germline depletion clusters both with 17α-E2 treatment, which requires an intact gonad for promoting male longevity^83,84^, as well as with castration (**Extended Data Figure 7b**). Since castration prevents the longevity benefit of 17α-E2 in mice, the physiological effects of 17α-E2 are expected to only partially overlap with castration (**Extended Data Figure 7b**, right). These intersections provide an important step toward identifying the molecular mechanisms that mediate the longevity benefits of germline depletion.

Our findings demonstrate that germline depletion can enhance longevity and somatic regeneration in a vertebrate model, with no significant tradeoff with somatic growth. The sex-specific phenotypes suggest that the effect is probably mediated by mechanisms other than the classical tradeoff between life-history traits^4,5^ - since a missing germline should similarly affect resource allocation in males and females (see a schematic model in **Figure 7c**). Next, we explored evolutionary conservation and possible effects on organismal healthspan.

### Evolutionary conservation with germline ablated worms

To investigate evolutionary conservation, we tested whether candidate genes that emerged from our single-cell transcriptomic data can mediate the lifespan extension in germline-depleted worms. We selected the CF1903 strain^85^, which harbors a temperature-sensitive mutation in *glp-1* (glp-1/Notch intracellular domain, *e2144*), an essential gene for germline maintenance in worms^85^. We confirmed that worm mutants display germline depletion and lifespan extension following a shift to the restrictive temperature (from 15 to 25°C)^85^. Interestingly, we observed a ‘neck-and-shoulder’ shape of the survival curve in *glp-1* mutants (**Figure 7d**, center, **Supplementary Table S4**), which is similar to our killifish data (**Figure 7b**).

We next selected several candidate genes that were identified in killifish (**Extended Data Figure 6d**), and perturbed these genes using RNAi in the *glp-1* mutants at the restrictive temperature. When compared to fertile controls (EV at 15°C), a similar longevity effect was observed in both sterile EV and *atf4* (transcription factor atf-4 homolog) RNAi (at 25°C). In contrast, *hsp1* (heat shock protein hsp-1) RNAi were shorter lived compared to the fertile controls, suggesting that loss of *hsp1* might be detrimental. Interestingly, the lifespan of sterile worms fed with *eef1a1* RNAi was not significantly longer compared to fertile worms (**Figure 7d**, center, **Supplementary Table S4**).

To further confirm the effect of *eef1a1*, we performed two additional comparisons. First, we observed no difference between fertile *eef1a1-* RNAi and EV-fed worms (at 15°C, **Extended Data Figure 7c**, left). Next, we directly compared the lifespan of sterile worms fed with either EV or *eef1a1* RNAi and validate that *eef1a1* RNAi-fed worms were indeed shorter lived (**Figures 7d**, right, **Extended Data Figure 7c**, right). These data suggest that fine-tuning the level of *eef1a1* is required for the lifespan extension of sterile worms.

### A distinct transcriptional response in old *dnd1*^Δ4^*^/^*^Δ4^ males

So far, we have characterized the transcriptome of young fish. To further investigate the effect of germline depletion on the aging process, we characterized the hepatic transcriptome of old WT or germline-depleted fish (25 weeks old, **Figure 8a**). Principle component analysis (PCA) indicated that germline depletion produced widespread transcriptional changes in the liver (**Extended Data Figure 8a**), with some linked to age (PC1), or to sex and genotype (PC2). The differential expression analysis in old fish revealed a total of 135 downregulated and 295 upregulated genes (FDR of 0.05, **Supplementary Table S8**). We next conducted gene set enrichment analysis (GSEA) using gene ontology (GO), combining datasets from both young and old fish.

**Figure 8:**
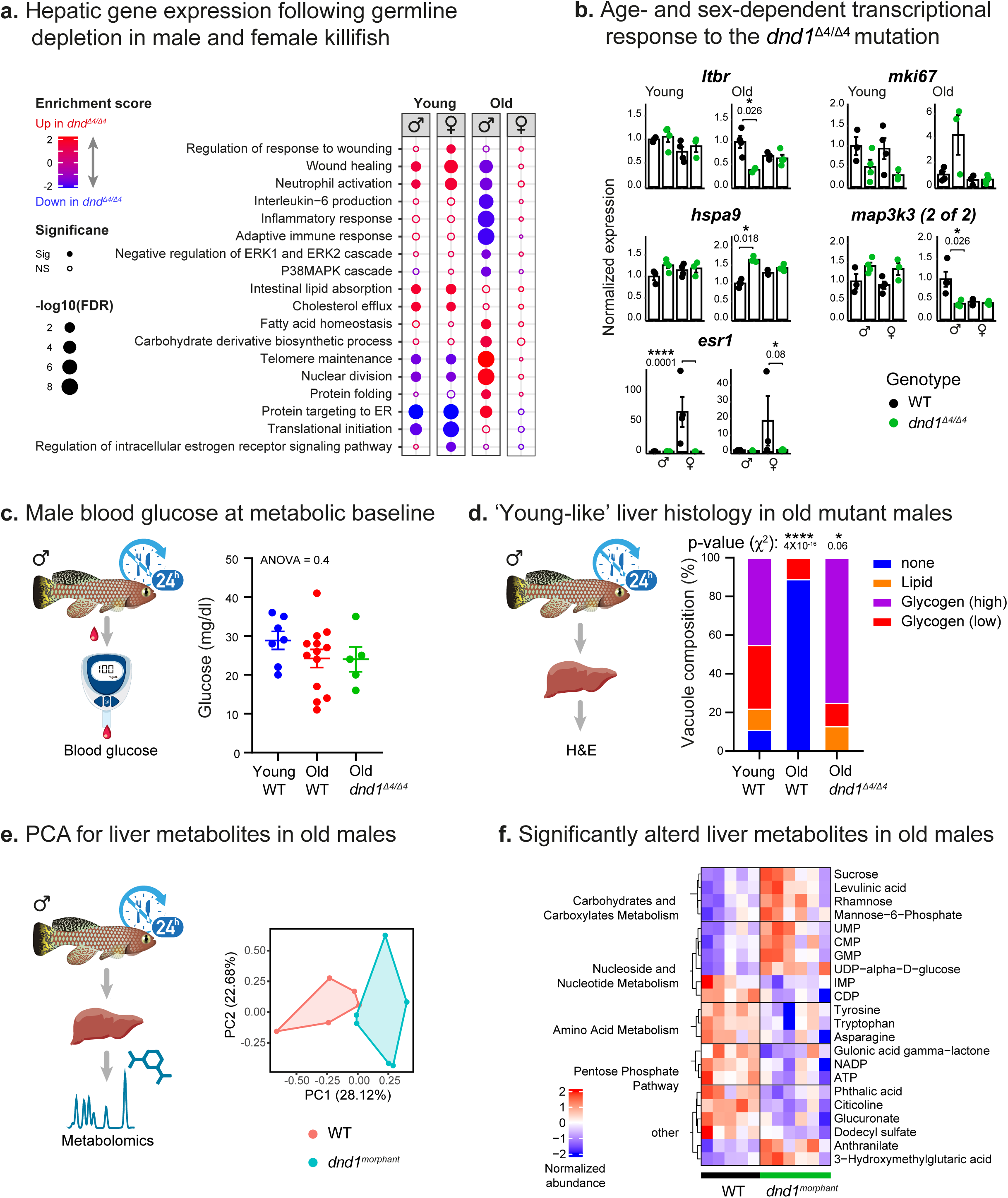
Germline depletion rejuvenates male hepatic metabolism. (**a**) Dot plot for functional enrichments (GO, FDR < 5%) using GSEA for differential gene expression between WT and *dnd1*^Δ4^*^/^*^Δ4^ mutant fish. Sex and age are indicated. (**b**) Relative expression (normalized to male WT) of driver genes from the pathways in (**a**). Error bars show mean□±□SEM with individual points from n = 3 - 4 individuals. Significance was calculated as part of the edgeR pipeline in differential expression analysis (generalized linear models and quasi-likelihood F-test), between the WT and *dnd1*^Δ4^*^/^*^Δ4^ mutant fish. (**c**) Blood glucose levels of the indicated experimental groups at metabolic baseline. n = 7 individuals for young WT, 13 for old WT and 5 for old *dnd1*^Δ4^*^/^*^Δ4^. Error bars indicate mean□± SEM. Significance was calculated using one-way ANOVA and exact p-value is indicated. (**d**) Quantification of liver vacuole composition in male fish of the indicated experimental condition at metabolic baseline (according to Figure 5c). Data presented as proportion of each vacuole-type. n ≥ 5 individuals for each experimental group. Significance was measured by two-sided χ^2^ test with the young WT proportion as the expected model and FDR correction. P-values are indicated. (**e**) PCA for hepatic metabolite levels of WT (red) or *dnd1^morphant^* (blue) old male fish. n ≥ 5 samples per condition. Each symbol represents an individual fish. (**f**) Heatmap showing metabolites that display a significantly altered response between old WT and germline-depleted male fish at metabolic baseline, significance was calculated using a two-sided Student’s t-test (p<0.05). Each square represents normalized relative abundance, from an average of n ≥ 5 fish.

While some enrichments in young fish were similar in males and females, several behaviors displayed a sex-specific pleiotropic pattern. For example, nuclear division is down regulated in young mutants of both sexes, and up regulated specifically in old male mutant (**Figure 8a**). Representative driver genes displaying sex-specific patterns in old age are also indicated (**Figure 8b**), including *mki67* (marker of proliferation ki-67) in ‘nuclear division’, *ltbr* (lymphotoxin beta receptor) in ‘inflammatory response’, and *hspa9* (heat shock protein family a member 9) in ‘protein folding’. We next set out to investigate which of these biological processes might be rejuvenated in old germline-depleted males.

### Rejuvenated metabolic functions in *dnd1*^Δ4^*^/^*^Δ4^ males

Our transcriptional analysis identified several pro-longevity pathways in germline-depleted males, including protein folding and carbohydrate metabolism (**Figures 6d, 8a**). Therefore, in addition to the irradiation stress described above, we decided to test other stressors. Furthermore, while *dnd1* expression is primarily restricted to the germline (**Extended Data Figure 8b**, top left), we cannot rule out a function for this gene beyond the germline. Therefore, to complement our genetic strategy, we used antisense morpholino injections against *dnd1*^7,65^ (see **Methods**). In agreement with prior studies^7,65^, injection of *dnd1* morpholino briefly and reversibly inhibits *dnd1* function during early embryogenesis (when the germline migrates into the gonad).

Morpholino injections were efficient in depleting the killifish germline (∼90% efficiency, see **Methods**), and were able to replicate our sex-specific findings following whole-body irradiation (given natural experimental variation, **Extended Data Figure 8b**, bottom). These data imply that the physiological effects are likely mediated by the depletion of the germline, and not via *dnd1* functions. We next tested other stressors, including proteostatic or oxidative stress (**Extended Data Figure 8c**, chloroquine or paraquat treatments, respectively). However, the regenerative potential was only modestly improved in germline-depleted males (particularly following proteostatic stress, **Extended Data Figure 8d**).

In agreement with the observed transcriptomic pattern (**Figure 8a**), quantification of proliferation using EdU demonstrated that germline depletion increases hepatic proliferation in old age (**Extended Data Figure 8e**). Another prediction from our RNA sequencing data is that germline depletion alters carbohydrate and lipid homeostasis, which naturally declines in old killifish^32^. While blood glucose content at metabolic baseline was not a good aging marker (**Figure 8c**, in agreement with recent fingings^86^), liver histology revealed that vacuoles, both lipid and glycogen type, are restored to a ‘young-like’ composition in old germline-depleted fish (**Figure 8d**).

To better assess the impact of germline depletion on the killifish fasting response, we applied metabolomics to livers isolated from WT and germline-depleted males at metabolic baseline^32^ (**Figure 8e, f, Supplementary Table S9**). All 140 unique metabolites passed our quality control (see **Methods**). Samples seem to segregate according to experimental conditions (PC1, **Figure 8e**), and differentially abundant metabolites were analyzed according to BioCyc^87^ and MetaboAnalyst^88^ (see **Methods**. **Figure 8f**). For example, a reduction in the pentose phosphate pathway (PPP) was observed in old germline-depleted fish, as well as altered nucleoside and nucleotide metabolism (i.e. increased GMP and UMP, **Figure 8f**). The PPP is an integral part of the mammalian fasting response^89^, and inhibition of the PPP promoted longevity in *C. elegans* by mimicking dietary restriction^90^. These trends partially overlap with our recent characterization of a youthful fasting response in killifish^32^.

Collectively, the transcriptomic, metabolomic, and histological data suggest that beyond the observed longevity phenotype, germline-depletion might also enhance male healthspan (such as metabolic functions). Ultimately, identifying the signals involved in mediating this phenotype could provide an exciting opportunity to uncouple the evolutionary link between reproduction and longevity.

## Discussion

A tradeoff between life-history traits is traditionally associated with reproduction, repair of somatic damage, growth, and lifespan^91^. However, the evidence that some of these tradeoffs can be experimentally uncoupled in invertebrate models, suggests the existence of alternative mechanisms for regulating life-history^92^. For example, following germline depletion in worms, the reproductive system can have a direct effect on hermaphrodite longevity via endocrine signals^8,85^. In contrast, complete removal of the gonad (both the germline and the somatic gonad), has no effect in worms. Nonetheless, the direct effect of specific germline manipulations on vertebrate life-history, particularly lifespan, has not yet been thoroughly evaluated^76^.

Here, we genetically perturb reproduction by manipulating the germline at discrete developmental stages. Our data suggest that as an alternative to the classic view, that any perturbation of reproduction can free up resources for somatic functions^93^, different forms of infertility produce different effects on life-history. Thus, the paradigms identified in worms might be conserved in a vertebrate model. It is worth mentioning that the observed longevity phenotypes in worms might be context-dependent. For instance, under favorable environmental conditions (e.g. moderate temperatures), the germline might not have a negative effect on killifish lifespan, and therefore might contribute to temperature-dependent variations in longevity^94^. Beyond lifespan, other downstream pathways are enhanced in the soma of germline-ablated worms, including proteostasis and energy/lipid metabolism^9^. Our data also suggests that metabolic functions might be rejuvenated (**Figure 8d-f**), and an additional mild effect was detected on proteostasis (**Extended Data Figure 8c-d**).

Chen et al.^7^ have recently proposed a trade-off between somatic and germline repair. Specifically, they have demonstrated that germline-depleted zebrafish can better regenerate their tail following irradiation. However, our data suggests that the regenerative potential of the tail could be independent of whether the germline is damaged or not (**Figure 5a, b**). Additionally, the sex-specific phenotypes seen in killifish following germline depletion further support the concept that some mechanisms might not depend on a classical tradeoff. However, it should be noted that regeneration of WT fish following irradiation was itself sex-dimorphic (**Figures 4d, Extended Data Figure 4d**). This could suggest that male fish might innately possess enhanced regenerative potential, as the tail plays an important role in mate recognition^95^. Additionally, the differences in regenerative potential between zebrafish and killifish might be attributed to innate physiological differences, such as the all-male sex reversal observed in germline-depleted zebrafish^7^.

What can mediate the observed sexual dimorphism? Plausible candidates are the sex-dimorphic circulating hormones (**Figure 3a**). Notably, both pituitary and sex hormones have previously been linked to vertebrate longevity, such as growth hormone or its receptor^96^ . However, as germline-depleted fish were of normal size (**Extended Data Figure 4a**), other circulating factors might be at play. In humans, exceptional longevity was recently shown to be correlated with the preservation of the integrity of the HPG axis^97^. Modified levels of steroid hormones have also been reported to be linked to longevity^98^, regeneration, metabolism, and immune functions (e.g. testosterone is an immunosuppressant^99^). Yet, it is worth mentioning that the pursuit to identify systemic regulators in germline-ablated worms has been ongoing for decades^8,19–22,85^.

Gonadal development, particularly gonadal size, depends on the presence of the germline in many species, from humans to worms^8^. Accordingly, following germline depletion, the gonad is affected and becomes hypoplastic, which could contribute to the observed phenotypes. Importantly, an “empty” vertebrate gonad can still retain a partial signaling ability^14,39,100–103^. To further investigate this, we applied several strategies. First, by comparing the hepatic transcriptional signature in germline-depleted fish to common longevity interventions in mice, we identify an overlap with 17αE2 treatment, which is thought to be naturally synthesized outside the gonad^104^. Similar to 17αE2-treated mice, germline-depleted killifish males display rejuvenated metabolic functions and reduced ‘inflammaging’ (**Figures 7, 8**), and the longevity effect of 17αE2 in mice is male-specific and depends on the presence of a gonad^79,84^.

Further comparison with gonadal/hormonal interventions in male mice (e.g. castration, estrogen receptor KO, hormone replacement) indicated that germline depletion still clusters with both 17α-E2 treatment and castration (**Extended Data Figure 7b**). Our recent longevity intervention in killifish, through indirect activation of AMPK (mitogen-activated protein kinase 1), was also male-specific and displayed a similar 17αE2 signature (which involved AMPK activation^80^). These data raise the possibility that part of the effect of germline depletion on lifespan is related to the AMPK pathway. These partial overlaps are an important step towards identifying downstream mechanisms, and support a possible involvement of the remaining somatic gonad.

Curiously, which sex undergoes lifespan extension following germline depletion varies across evolution. In androdiecious worm species (with hermaphrodites and males, such as *c. elegans*) germline depletion causes lifespan extension primarily in hermaphrodites^8^. However, in gonochoristic worm species with true males and females, such as *C. remanei*, it seems to affect mostly males^105^. Because of this complexity, to functionally test candidate genes in worms, we chose differentially expressed genes from killifish regardless of their expression pattern, including upregulated in males and downregulated in females (*hsp1a1*), up in both sexes (*atf4*), or down in males and up in females (*eef1a1*). We chose RNAi and not overexpression, as overexpression requires gene-specific information, such as at which tissues or developmental stages each gene is normally expressed in worms. Our data suggests that the fine-tuning of protein translation through *eef1a1* is required for the longevity effect of germline-depleted worms (**Figures 7d**, **Extended Data Figure 7c**). Accordingly, while *eef1a* expression is downregulated in germline-depleted long-lived male killifish gonads (**Extended Data Figure 6d**), feeding *eef1a1* RNAi to germline-depleted *C. elegans* hermaphrodites prevents lifespan extensions.

The timing of germline depletion might be important in mediating the physiological effect. For example, while early germline depletion in *Drosophila* shortened the female lifespan (with a modest extension of the male lifespan)^106^, exclusive elimination of germ cells in late development extended the lifespan of both sexes^107^. The intriguing correlative data in humans and several other farm and laboratory animals demonstrated that removing the entire gonad (e.g. Korean eunuchs, sheep castration) can have profound longevity effects in males (with complex phenotypes following females ovariectomy^10–17)^. However, the timing differs from the outcome in *Drosophila*, as men castrated before sexual maturation lived longer than if the castration occurred after sexual maturity^10^. Therefore, it will be important to develop genetic tools to directly compare castration and germline depletion, or to induce germline depletion at specific time points.

Finally, germline depletion in killifish primarily affects maximal lifespan (**Figure 7b**). Thus, suggesting a ‘temporal window’ in which longevity benefits are provided primarily during late-life. These findings are in contrast to effects on median lifespan produced by metabolic interventions, such as in the worm *eat-2* mutants^108^ (nicotinic acetylcholine receptor subunit) and the killifish *aprt* mutants^32^ (adenine phosphoribosyltransferase). In conclusion, the means by which the germline manipulates organismal physiology could explain how the ratio between age at maturity and organismal lifespan is evolutionarily conserved across so many vertebrate species.

## Methods

### African turquoise killifish strain, husbandry, and maintenance

All turquoise killifish care and uses were approved by the Subcommittee on Research Animal Care at the Hebrew University of Jerusalem (IACUC protocols #NS-18-15397-2 and #NS-22-16915-3).The African turquoise killifish (GRZ strain) was housed as previously described^31,33^. Fish were grown at 28°C in a central filtration recirculating system with a 12 h light/dark cycle at the Hebrew University of Jerusalem (Aquazone ltd, Israel). Fish were fed with live Artemia until the age of 2 weeks (#109448, Primo), and starting week 3, fish were fed three times a day on weekdays (and once a day on weekends), with GEMMA Micro 500 Fish Diet (Skretting Zebrafish, USA), supplemented with Artemia once a day. Under these conditions, the median killifish lifespan was approximately 4-6 months. All genetic models (described below) were maintained as heterozygous and propagated by crossing with wild-type fish.

### CRISPR/Cas9 target prediction and guide RNA (gRNA) synthesis

CRISPR/Cas9 genome-editing protocols were performed as described previously^31^. Briefly, conserved regions upstream of functional or active protein domains were selected for targeting the selected genes. gRNA target sites were identified using CHOPCHOP (https://chopchop.rc.fas.harvard.edu/)^109^. When needed, the first base-pair was changed to a ‘G’ to comply with the T7 promoter. Design of variable oligonucleotides, and hybridization with a universal reverse oligonucleotide was performed according to^31^, and the resulting products were used as a template for *in vitro* transcription. gRNAs were *in vitro* transcribed and purified using a quarter reaction of TranscriptAid T7 High Yield Transcription Kit (Thermo Scientific #K0441), according to the manufacturer’s protocol. All primer and gRNA sequences used to generate mutant lines can be found in **Supplementary Table S10**.

### Production of Cas9 mRNA

Experiments were performed as described previously^30,31^. The pCS2-nCas9n expression vector was used to produce *Cas9* mRNA (Addgene, #47929)^110^. Capped and polyadenylated *Cas9* mRNA was *in vitro* transcribed and purified using the mMESSAGE mMACHINE SP6 ULTRA (ThermoFisher #AM1340).

### Microinjection of turquoise killifish embryos and generation of mutant fish using CRISPR/Cas9

Microinjection of turquoise killifish embryos was performed according to ^31^. Briefly, nCas9n-encoding mRNA (300 ng/μL) and gRNA (30 ng/μL) were mixed with phenol-red (P0290, Sigma-Aldrich) and co-injected into one-cell stage fish embryos. Sanger DNA sequencing was used for detecting successful germline transmission on F1 embryos. Fish with desired alleles were outcrossed further to minimize potential off-target effects and maintained as stable lines by genotyping using the KASP genotyping platform (Biosearch Technologies) with custom made primers. All primers used in generating the mutations can be found in **Supplementary Table S8**.

### Microinjection of turquoise killifish embryos with morpholino

When indicated, as an alternative approach for mutating *dnd1,* progenitor germline stem-cells were depleted using morpholino injection, as previously performed for zebrafish^7,65^. Briefly, fertilized eggs, at the 1-4 cell stage, were injected with a solution of a custom made antisense morpholino against the *dnd1* gene (100 μM, GAC GGT CTT CCT CTC CAT GTC, Gene Tools, USA) diluted in 0.2 M KCl. 1.5% Rhodamine B isothiocyanate–Dextran (Sigma-Aldrich, #R8881) was also added to the injection solution to ease visual confirmation of successful injection by fluorescence signal. Success of germline depletion following morpholino injection was visually confirmed by dissection, with ∼90% (44/49) success rate. Fish that were discovered to have a germline were censored from the experiments (see **Supplementary Table S4**).

### METHOD DETAILS

### Single cells RNA-seq

#### Dissociation of gonads for scRNA sequencing

For wild-type fish, three gonads were combined from either one-month old males or females, while for *dnd1*^Δ4^*^/^*^Δ4^ mutants, five fish were pooled per group (as the gonads are very small). Gonads were dissected from each fish according to^111^ and kept on ice in full medium (L15, 1% penicillin-streptomycin, 50 μg/μl gentamicin, 15% FBS). When all dissections were complete, organs were washed once with L15, and incubated in digestion media (400 μl 0.25% trypsin) at 28°C for 2 h with periodical passage through a glass Pasteur pipette to mechanically aid dissociation. This was followed by the addition of 800 μl full medium to stop enzymatic digestion. Dissociated cells were passed through a 100 μm cell strainer prior to FACS analysis. For live/dead sorting by FACS, cells were incubated with propidium-iodide (PI, 1 μg/ml) for 5 minutes, and live cells were sorted according to PI intensity using an Aria III Sorter (BD) with the BD FACSDiva 8.0.1 software. Live cells were used for downstream single cell RNA sequencing. ∼70% of cells in the sample were alive and downstream RNA-seq did not detect signatures of dead cells. Gating strategy can be found in **Supplementary Figure S1**. Single cell RNA-seq library preparation and microfluidic operations were performed using the Fluigent microfluidic pump system in a microfluidic device, that was fabricated by soft lithography following standard InDrop protocols^45,112^ with some adaptations^44,113^. A detailed protocol is available in the **Supplementary Methods** section.

#### Sequencing and alignment

The reads were split into barcode-specific files for mapping. Read 1 was used to obtain the sample barcode and UMI sequences, and read 2 was aligned to the reference genome. The alignment was performed using RSEM (v1.3.1)^114^ and Bowtie2 (v2.4.5)^115^ with default parameters and the single-cell mode (--single-cell-prior) using the killifish reference genome (Nfu_20140520^34,35^).

#### scRNA sequencing analysis

##### Quantification and statistical analysis

For scRNA-seq data analysis the transcript per million (TPM) values were used, as an input for Scanpy (v1.9.1)^116^ pipeline on Python 3.9.7. Cells with between 200 and 4000 unique genes and with less than 20% mitochondrial genes were retained for further analysis. Identification of 1000 highly variable genes was performed with the following parameters: flavor=“seurat_v3”, n_top_genes=1000. A global-scaling normalization was performed on the filtered dataset using “normalize_total” with a scale factor of 10,000, “log1p” with default parameters, and “scale” with max value of 10.

Cell-to-cell variation in gene expression driven by batch, cell alignment rate, and the number of detected molecules were regressed out, and linear transformation was applied. Harmony (v0.1.7)^46^ correction was used to verify that there is no batch effect in our data. PCA was performed on the scaled data. K-nearest neighbors were calculated using “sc.pp.neighbors” with 20 neighbors and 15-20 PCs. Then, the graph was embedded in two dimensions using UMAP with min_dist=0.4-0.5 and the cells were clustered the using the Leiden algorithm with a resolution of 0.2-0.7.

WT cells (males and females) were analyzed to identify germ cell clusters using known markers (*ddx4*, *dmc1*, *tekt1,* and *zp4*). Germ cells were then filtered out and analyzed separately. Male (WT and *dnd1*) and female (WT and *dnd1*) cells were analyzed separately, and all somatic cell types were identified. Finally, clusters that displayed an overlap between the genotypes, were selected for differential gene expression analysis. In the germline, identified erythroid and female cells were filtered out. Male germ cells were analyzed using pseudotime^117^ with a root cell in cluster 2 as well as a clustering analysis that was similar to the somatic cell analysis.

##### Differential expression analysis

Differential gene expression was performed using “sc.tl.rank_genes_groups” with the following parameters: corr_method=“bonferroni”, method=“wilcoxon”, pts=True, tie_correct=True. The log fold change between two groups was calculated according to Seurat^118^ (log2 of mean (A + 1) / mean (B + 1)).

##### Gene Ontology Enrichment Analysis

Enriched Gene Ontology (GO) terms were identified using GO (clusterProfiler v3.18.1)^119^, with the threshold of p-value < 0.1 and FC > 1.5. GO terms were based on human GO annotations from org.Hs.eg.db (v3.13.0)^120^ and AnnotationDbi (v1.54.1)^121^.

##### Identification of marker genes

The differential expression between each cluster and the remaining clusters was calculated. Cell types were identified using the top differentially expressed genes, and their overlap with classical cell-type-specific markers^49–52^. Full lists of the marker genes are available in **Supplementary Tables S1, S2, S3, S5, and S6**. The differential expression analysis between WT and *dnd1* mutants (within a specific cell type) was then used to perform enrichment analysis using GO with FDR < 0.1 and log2-fold-change of 1.5 using clusterProfiler (v4.0.5)^119^ and org.Hs.eg.db (v3.13.0)^120^ R packages (v4.1.1). Full lists of differentially express genes and GO pathways are available in **Supplementary Table S7**.

##### Evolutionary comparison with single-cell data from zebrafish gonads

We directly compared our identified cell types with male and female zebrafish single-cell data: Qian et al. 2022^43^ for male germline comparison. The top five differentially expressed genes of each cluster in zebrafish were selected, and their expression in the killifish clusters was examined.

### Bulk RNA-seq of the livers

#### Preparation of livers for RNA sequencing

Fish were grouped by sex (males or females), genotype (WT or *dnd1*^Δ4^*^/^*^Δ4^), and age (5- or 25-week-old). For each experimental group 3-4 fish were used. Fish were euthanized in 500 mg/L of Tricaine (Sigma-Aldrich, A5040) in system water. Livers were collected and flash frozen in liquid nitrogen and kept in -80°C until use. Tissues were homogenized by metal beads in 300 μl of TRI reagent (Sigma-Aldrich, T9424), using a TissueLyzer LT (QIAGEN, #85600) with a dedicated adaptor (QIAGEN, #69980). RNA purification was performed using the Direct-zol RNA Miniprep kit (Zymo, R2052) according to the manufacturer’s instructions. RNA concentration and quality were determined by using an Agilent 2100 bioanalyser (Agilent Technologies).

#### RNA-seq library preparation

Library preparation was performed using KAPA Stranded mRNA-Seq Kit (ROCHE-07962193001) according to the recommended protocols. Library quantity and pooling were measured by Qubit (dsDNA HS, Q32854), with size selection at 4% agarose gel. Library quality was measured by Tape Station (HS, 5067-5584). Libraries were sequenced by NextSeq 2000 P3, 50 cycles, 70 bp single-end (Illumina, 20046810) with ∼35 million reads per sample.

#### RNA sequencing analysis

Quality control and adapter trimming of the fastq sequence files were performed with FastQC (v0.11.8)^122^, multiQC (v1.12)^123^, fastx-toolkits (v0.0.13), Trim Galore! (v0.6.4)^124^, and Cutadapt (v3.4) ^125^. Options were set to remove Illumina TruSeq adapters and end sequences to retain high-quality bases with *phred* score > 20 and a remaining length > 20 bp. Successful processing was verified by re-running FastQC. Reads were mapped and quantified to the killifish genome Nfu_20140520^34,35^ using STAR 2.7.6a^126^. Differential gene expression as a function of genotype was performed using the edgeR package (v3.32.1)^127,128^. The ComBat_seq function from the sva package (v3.42.0)^129^ was used for batch correction.

#### Gene Ontology Enrichment Analysis

Enriched Gene Ontology (GO) terms associated with transcripts levels (from WT versus *dnd1*) were identified using Gene Set Enrichment Analysis (GSEA) implemented in R package clusterProfiler (v3.18.1)^119^. All the transcripts were ranked and sorted in descending order based on multiplication of log2 transformed fold change and -log10(FDR). Note that due to random seeding effect in GSEA, the exact p-value and rank of the enriched terms may differ for each run. These random seeds did not qualitatively affect the enrichment analyses. GO terms were based on human GO annotations from org.Hs.eg.db (v3.13.0)^120^ and AnnotationDbi (v1.54.1)^121^. Heatmap visualization was perform using ComplexHeatmap (v2.8.0)^130^ with hierarchical clustering using Pearson correlation.

#### Principal component analysis (PCA)

Standardized log2 transformed normalized count per million (CPM) were used as input for PCA. PCA was performed using autoplot function implemented in R package ggfortify (v0.4.12) and plotted using ggplot2 (v3.3.5).

#### Comparing transcriptional longevity interventions between killifish and mice

The transcriptional signature of 8 longevity interventions in mouse liver were previously published^75^. The transcriptional signature of the *dnd1* mutation in killifish was obtained by comparing young WT and *dnd1* mutants, either male or female fish. The same enrichment strategy described in^75^ was applied, using GSEA and the msigdbr R package (v7.5.1).The data are presented in a heatmap of normalized enrichment score (NES)using ComplexHeatmap (v2.8.0)^130^ with hierarchical clustering using Pearson correlation.

#### Comparing transcriptional gonadal/hormonal interventions between killifish and mice

To compare the transcriptional signature of germline depletion in fish to other common gonadal/hormonal interventions in male mice (e.g. castration, estrogen receptor KO, hormone replacement) ^75,81,82^ we used a ranking list derived from the log2(FC) between intervention and control groups for enrichment analysis, as described above. The pathways were extracted from the msigdbr R package (v7.5.1), including biological process and molecular function (GO), KEGG, Reactome, and WikiPathways according to org.Mm.eg.db (v 3.14.0)^131^ annotation. The data are presented in a heatmap of normalized enrichment score (NES).

### Survival, growth, and metabolic assays

#### Lifespan measurements, killifish

Constant housing parameters are very important for reproducible lifespan experiments^31,132^. After hatching, fish were raised with the following density control: up to 30 fish in a 1-liter tank for week 1, 5 fish in a 3-liter tank for weeks 2-4. From this point onwards, at the age of 4 weeks, adult fish were genotyped and housed individually in a 1-liter tank for the rest of their life. Plastic plants were added for enrichment. Both male and female fish were used for lifespan experiments and were treated identically. Fish mortality was documented daily starting at week 4. Lifespan analyses were performed using GraphPad Prism for all survival curves with a Kaplan-Meier estimator. A log rank test was used to examine any significant differences between the overall survival curves of different experimental groups. Median and maximal (90^th^ percentile) lifespan were calculated (**Supplementary Table S4**). Boschloo’s test for the 50th, 75th and 90th percentile was performed using the on-line tool OASIS 2^78^. Quantile-Quantile plots were calculated using R and plotted in GraphPad Prism.

#### Lifespan measurements using temperature sensitive sterile nematodes

Experiments were performed according to^133^. Briefly, synchronized worm populations were grown on Nematode Growth Medium (NGM) plates supplemented with 100 µg/ml ampicillin and seeded with E. Coli bacteria (strain HT115) that harbor the indicated RNAi plasmids. All the RNAi used in the study were taken from the “VIDAL” RNAi library^134^. To induce the expression of dsRNAi, the seeded plates were supplemented with 100 mM isopropyl β-d-1-thiogalactopyranoside (IPTG; final concentration of 4mM). To induce the expression of the dsRNA. To test whether the genes of interest are required for lifespan extension by germline ablation we used the CF1903 worm strain^85^. These animals carry a temperature-sensitive mutation in their *glp-1* gene which renders them sterile when grown at 25°C during larval development. However, these animals are fertile when developed at 15°C. This mutation leads to a significant extension in the lifespan of sterile worms as compared to the control, fertile worms. Two control groups were included in each experiment, both grown on bacteria that harbor the empty RNAi plasmid (EV). While worms of one group developed at 15°C and thus, exhibited normal reproduction profile and natural lifespan, animals of the other group developed at 25°C and therefore, had no germline and were long-lived. All RNAi-treated worms developed at 25°C. At day 1 of adulthood worms of all groups were transferred onto 60 mm NGM plates (8-10 plates for each treatment, 12 animals per plate) and incubated subsequently at 20°C. Living animals were scored daily until the last worm of all treatments died. Lifespan analyses were performed using GraphPad Prism for all survival curves with a Kaplan-Meier estimator. A log-rank test was used to assess the difference between pairs of lifespan curves and then all p-values were corrected using FDR, when required.

#### Growth measurements

For growth assays, a single fish was housed in a 1-liter tank from week 2. Both sexes were measured by imaging at the indicated timepoints with a Canon Digital camera EOS 250D. To limit vertical movement during imaging, fish were placed in 3 cm deep water in a tank, and images were taken from the top. A ruler was included in each image to provide an accurate scale. Body standard length was measured from the tip of the snout to the posterior end of the last vertebra. The length was then calculated by Matlab (R2021a), by converting pixel number to centimeters using the reference ruler.

#### Fertility Analysis

Fish fertility was evaluated as described previously^30,32^. Briefly, 3-6 independent age-matched pairs of fish (one male, one female) of the indicated genotypes were placed in the same tank. All breeding pairs were allowed to continuously breed on sand trays, and embryos were collected and counted on a weekly basis for 4 weeks. Results were expressed as the number of eggs per couple per week of egg-lay. Significance for each genotype (compared to the WT) was calculated using one-way ANOVA with a Dunnet post-hoc in Prism (GraphPad).

#### Blood glucose measurements

5-week-old (young) and 12-week (old) fish were fasted for 24 hours following morning feeding. Caudal fin was removed, and blood was collected directly to an Accu-check Instant blood glucose meter (Roche). Significance was calculated using one-way ANOVA.

### Polar metabolic profiling

#### Organ collection

12-week-old fish were grouped by genotype (wild-type or *dnd1*^Δ4^*^/^*^Δ4^), and fasted for 24 hours following the morning feeding. 5-6 fish were collected for each group. Fish were euthanized in 500 mg/L of Tricaine (Sigma-Aldrich, A5040) in system water. Dissections were carried out under a stereo binocular (Leica S9D) at room temperature. Livers were collected, flash frozen in liquid nitrogen, kept in -80°C until use.

#### Sample preparation

Polar metabolites were extracted at the Life Sciences Core Facilities, Metabolic Profiling Unit (Weizmann Institute of Science), as previously described^135,136^ with some modifications. Briefly, liver samples were lyophilized, ground to powder, and mixed with 1 mL of a pre-cooled (20°C) homogenous methanol:methyltert-butyl-ether (MTBE) 1:3 (v/v) mixture. The tubes were vortexed and then sonicated for 30 min in an ice-cold Transsonic 460/H sonication bath (Elma) at 35 kHz (taken for a brief vortex every 10 min). Then, UPLC-grade water (DDW): methanol (3:1, v/v) solution (0.5 mL) containing internal polar metabolite standards (C13 and N15 labeled amino acids standard mix (1:500), Sigma, 767964) were added to the tubes. Following 5 min centrifugation at maximum speed, the upper lipid phase was removed, and the polar phase was re-extracted as described above, with 0.5 mL of MTBE, transferred to new tube, lyophilized, and stored at -80°C until analysis.

#### LC-MS for polar metabolite analysis

The lyophilized pellets were dissolved using 150 mL DDW-methanol (1:1), centrifuged twice (at maximum speed) to remove possible precipitants, and injected into the LC-MS system. Metabolic profiling of the polar phase was performed as previously described^137^ with minor modifications described below. Briefly, analysis was performed using an Acquity I class UPLC System combined with a mass spectrometer (Thermo Exactive Plus Orbitrap) operated in a negative ionization mode. The LC separation was performed using the SeQuant Zic-pHilic (150 mm 3 2.1 mm) with the SeQuant guard column (20 mm 3 2.1 mm) (Merck). Mobile Phase B was acetonitrile; Mobile Phase A consisted of 20 mM ammonium carbonate with 0.1% ammonia hydroxide in DDW:acetonitrile (80:20, v/v). The flow rate was kept at 200 mL/min and gradient as follows: 0-2 min 75% of B, 17 min 12.5% of B, 17.1 min 25% of B, 19 min 25% of B, 19.1 min 75% of B, 23 min 75% of B.

#### Polar metabolites identification and quantification

The data processing was performed using TraceFinder 4 software (Thermo Fisher), and compounds were identified by accurate mass, retention time, isotope pattern, and fragments and verified using in-house mass spectra library.

#### Metabolomic analysis

Samples were normalized by internal standard and total protein. Metabolites were omitted if they were detected in less than 70% of the samples (20/28 samples), with a final list of 140 metabolites. Values were log2 transformed and normalized by the average of each metabolite. Hierarchical clustering was based on Pearson correlation. Significance was calculated using a two-sided unpaired t-test between the WT and *dnd1* mutant, and assignment into specific metabolic pathways was done according to BioCyc^87^ and MetaboAnalyst^88^. Metabolites that are part of several pathways were assigned to a single pathway. Significant metabolites (p-value < 0.05) are presented in a heatmap (**Supplementary Table S9**)

### Histology

#### Hematoxylin and eosin

Tissue samples were processed as described previously^29,30,32,33,35,36,111,138–145^. Briefly, paraffin sections from one-month old fish of the indicated sex and genotype were used. 5-10 μm sections were stained with Hematoxylin and Eosin, and examined by microscopy. Stages of egg and sperm development were identified according to^146^. For each liver, vacuoles were quantified from three different images, and scored according to^66^ and the scale shown in **Figure 5c**. Adipose tissues were identified according to^67,68,147^ and scored according to the scale shown in **Figure 5c**.

#### Florescent in-situ hybridization (FISH)

Split initiator hybridization probes for florescent in-situ hybridization chain reaction (HCR v.3.0) were designed and manufactured by Molecular Instruments for the following genes: *lhr* (XM_015944467.1), *fshr* (XM_015962932.1), *amh* (XM_015977279.1), *star* (XM_015963550.1), ddx4 (XM_015957842.1), *ptgds* (XM_015955369.1), *col1a1* (XM_015968981.1), *dmc1* (XM_015963686.1), *tekt1* (XM_015977050.1), *zp4* (XM_015956918.1), *cyp19a1* (XM_015964243.1), *eef1a1* (XM_015974153.1), and *hsp1a1* (XM_015941026.1). 10 µm paraffin slices (as described above), were baked for 1 h at 60°C, and FISH was performed according to the manufacturer’s guidelines. Briefly, following rehydration, slides were boiled for 15 minutes in 0.01 M citrate buffer (Sigma-Aldritch #C8532) and permeabilized with 20 µg/ml proteinase K (A&A Biotechnology #1019-20-5) in PBS for 15 minutes at 37°C, before being washed with PBS. Slides were pre-hybridized in hybridization buffer provided by the manufacturer (Molecular Instruments), and then hybridized with the indicated probe (20 nM, in hybridization buffer), at 37°C over-night. After washing, the signal was amplified with either custom-made green (488) or red (546) fluorophores at room temperature overnight. Slides were washed with 5XSCCT, and autofluorescence was quenched using Trueview autofluorescence quenching kit (Vector Labs #SP8500) according to the manufacturer’s protocol. Slides were mounted with Vectashield containing DAPI (Vector Labs #30326), and images were collected with an Olympus FV-1200 confocal microscope and processed in imageJ^148^.

#### Immunohistochemistry

One-month old fish were euthanized with MS222 (500 mg/l) for 10 min at room temperature. The upper jaw and head were dissected and fixed in 4% PFA in PBS overnight at 4°C, followed by decalcification in EDTA (Avanator Performance materials, #8993-01, 0.5M, pH 8) for 7 days at room temperature. The heads were then immersed in an OCT-glucose solution (20% sucrose (Bio-Lab #001922059100), 30% OCT (Scigen Scientific Gardena #4586), 50% PBS), overnight at 4°C, followed by embedding in OCT and freezing in liquid nitrogen. Serial 20 µm sagittal sections were cut on a cryostat, airdried, and stored at -80°C. For immunostaining, slides were washed in PBS, and permeabilized for 15 minutes in 0.25% Triton (Avanator Performance materials #X198-07), 1% BSA (Sigma-Aldritch #A7906 in PBS), followed by blocking (Dako #X0909) for 10 minutes, and incubation with primary antibodies overnight. The following primary antibody was used: anti-rabbit anti-carp LH antibody (1:100), a generous gift from Prof. Berta Levavi-Sivan^149^ and a donkey anti rabbit Alexa Fluor 549 secondary antibody (Abcam #150064, 1:500), for 1 h at room temperature. After several washes, autofluorescence was quenched using Trueview autofluorescence quenching kit (Vector Labs #SP8500) and mounted with Vectashield containing DAPI (Vector Labs #30326). Samples were imaged with an Olympus FV-1200 confocal microscope and processed in imageJ^148^.

#### Assessment of cell proliferation by EdU staining

12-week-old fish were grouped by experimental condition (wild-type or *dnd1^morphant^*) with additional 5-week-old WT fish. Fish were injected with 50 mg/kg fish weight of 5-Ethynyl-deoxyuridine (EdU) for 2 consecutive days, and then fasted for 24 hours following the morning feeding (to match the conditions of the RNAseq and Metabolomics). Liver samples from 5-6 fish from each group were dehydrated and paraffin sections were taken as described above. EdU staining was preformed using a ClickTech EdU Cell Proliferation Kit (BaseClick #BCK-EdU488IM100) according to the manufacturer instructions and mounted with Vectashield containing DAPI (Vector Labs #30326). Samples were imaged with a fully motorized Olympus IX23 microscope with an Olympus DP28 camera. At least 3 images of each liver were taken. Proliferation was calculated by dividing EdU^+^ cells by overall nuclear numbers using ImageJ^148^.

### Injection and ectopic over-expression of plasmids via electroporation

#### Cloning of killifish cDNAs

Total RNA was isolated from the lysed brain and gonads using the Direct-zol RNA miniprep kit. (Zymo research #R2052), and the Verso cDNA Synthesis Kit (Thermo scientific #AB1453A) was used to prepare cDNA with random primers according to the manufacturer’s protocol. cDNA for the *lhb* and *fshb* hormones was amplified using Platinum SuperFi II DNA Polymerase (Invitrogen, #12361010). Primer sequences are available in **Supplementary Table S8**. PCR products were purified (QIAquick PCR purification kit, Qiagen #28104). The sequence-verified ORFs were cloned using GIBSON (NEB, #E2611L) into the pLV-EGFP plasmid, which was modified such that each hormone is tagged with a GFP, separated by the T2A self-cleaving peptide^150^. Plasmids and corresponding annotated maps are available via Addgene (#194356, #194357)

#### In-vivo electroporation

The electroporation protocol was adapted from^38^. Briefly, fish were sedated in MS222 (200 mg/l), and 3-5 µl plasmid solution (Plasmid concentration 100-200 ng/µl) containing Phenol Red for visualization (0.1% Sigma #P0290) was injected intramuscularly using a Nanofil syringe (WPI, #NANOFIL). Fish were then electroporated using 7 mm tweezer electrodes (NEP GENE #CUY650P10) with the ECM 830 generator. (BTX #45-0661). Electrodes were coated with wet cotton to minimize the risk to the fish. Fish were electroporated with 6 pulses of 28 V for 60 ms each with an interval of 1 s between each pulse. Images were recorded by a Leica MC190HD camera mounted on a Leica M156FC microscope.

### Tail regeneration following irradiation and pharmacological treatments

#### Irradiation procedure

Irradiation, regeneration, and terminal deoxynucleotidyl transferase dUTP nick end labeling (TUNEL) protocols were adapted from previous reports^7,151^. Briefly, one-month-old-fish (young) or 5-month-old fish (old) of the indicated genotypes were exposed to a sublethal dosage of X-ray irradiation (20 Gy) using the XRad 320 irradiator (Precision X-ray) at the Weizmann Institute of Science or at the Hebrew University. To this end, fish were sedated with MS222 (200 mg/l), placed in a plastic tray with separate wells, and irradiated until the desired dosage of irradiation was reached (approximately 6 minutes). For tail-specific irradiation, fish were held by their tail to minimize movement, and an adjustable collimator was attached to the irradiator in order to limit the X-ray beam. Radiation outside the beam was validated by dosimeter. Fish were then transferred to separate tanks to recover. Regeneration assays were performed three days post irradiation.

#### Drug treatment

Fish were drug-treated in system water for six days post amputation. Water was changed every three days. Fish were fed shortly before water changes. Fish were treated with 4mM of chloroquine (Sigma-Aldrich, #C6628) or 1 mg/ml paraquat (Sigma-Aldrich, #856177).

#### Regeneration assay

Fish were sedated, placed on a damp Kim wipe, and approximately 50% of the tail was amputated with one clean cut using a razor blade. Images were collected with a Leica MC190HD camera mounted on a Leica M156FC microscope, before and after amputation, and then every 3 days for 18 days. The length of the tail, from the base of the tail to the posterior regenerating tip, was measured at three different points, and the values were averaged. Regrowth was calculated as the percentage of 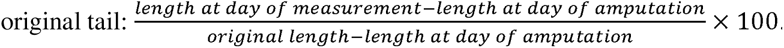.

#### TUNEL staining

Seven days post irradiation, the fish were euthanized with MS222 (500 mg/L), fixed for 72 h in 4% PFA at 4°C, and embedded in paraffin as described above. Tissue sections (10 µm) were stained using a TUNEL kit (Promega, #G3250) according to the manufacturer’s protocol and the samples were imaged with an Olympus FV-1200 confocal microscope and processed in ImageJ.

### Three-dimensional genome architecture (Hi-C)

#### Generation of fibroblast cultures from killifish tail fins

Fibroblast cells were prepared and cultured as described previously^32^. Briefly, following sedation, a 2–3 mm section of tissue was trimmed from the tail fin using a sterile razor blade, and disinfected for 10 min with a 25 ppm iodine solution (PVP-I, Holland Moran 229471000) in DPBS (Biological Industries). After rinsing with DPBS, tissue samples were incubated for 2 h with 1 ml of an antibiotic solution containing Gentamicin (50 µg/ml Gibco) and Primocin^TM^ (50 µg/ml, InvivoGen) in DPBS at room temperature. The samples were then transferred to an enzymatic digestion buffer (200 µl, in a 24-well plate) containing Dispase II (2 mg/mL, Sigma Aldrich) and Collagenase Type P (0.4 mg/mL, Merck Millipore) in Leibovitz’s L-15 Medium (Gibco), and were dissociated mechanically with a sterile pair of scissors. The digested tissue was mixed with 400 µl of complete Leibovitz’s L-15 growth medium (Gibco), supplemented with Fetal Bovine Serum (15% FBS, Gibco), penicillin/streptomycin (50 U/ml, Gibco), Gentamicin (Gibco, 50 µg/ml) and Primocin^TM^ (50 µg/ml, InvivoGen). For the first 7 days, cells were washed daily with fresh media before adding new media. The cells were incubated at 28°C in a humidified incubator (Binder, Thermo Scientific) with normal air composition until a stable cell-line was generated.

#### Generating Hi-C libraries

The Hi-C procedure used to map 3D genomic interactions in killifish was adapted from previous studies^152^, and detailed in the **Supplementary Method**s section.

#### Hi-C data processing and assembly scaffolding

As an initial step, Hi-C sequencing reads were processed against the previously published *Nothobranchius furzeri* genome assembly (GCA_001465895.2_Nfu_20140520) using the Juicer pipeline^153^ for analyzing Hi-C datasets (version 1.8.9 of Juicer Tools). The resulting Hi-C matrices were then used as input to the 3D DNA pipeline for automated scaffolding. Manual correction of obvious assembly and scaffolding errors was carried out using Juicebox^153^. After finalizing the scaffolding, Hi-C reads were reprocessed against the new assembly using the Juicer pipeline. The Hi-C assembly was aligned to the published genome (GCA_001465895.2_Nfu_20140520) by using LAST (https://gitlab.com/mcfrith/last/-/blob/main/doc/last-papers.rst).

### Statistics and Reproducibility

The number of biological replicates and statistical tests for each experiment are present in the corresponding figure legend. No statistical methods were used to pre-determine sample size but our sample sizes are similar to those reported in previous publications^7,32,38,133^. All fish were randomly assigned to each experimental group, while controlling for age, sex, and genotype. Blinding was used for all quantifications, including histology assessments and tail growth measurements. For lifespan and regeneration studies, animals that have died an unnatural death or immediately following radiation were excluded. *dnd1* mutants/morphants that displayed germline in their gonads were also excluded. These criteria were pre-established. Censored animals are described in **Supplementary Table S4**. Otherwise, no fish were excluded. Experiments were repeated independently with comparable findings. Experiments involving smFISH, immunostaining, TUNEL, liver histology, blood glucose, proliferation, and regeneration following stressor other than irradiation and of morphants were replicated twice. Experiments involving fertility, growth, gonad histology, regeneration following irradiation of genetic mutants and tail only irradiation and worm lifespan were replicated three times. This excludes the omics datasets and fish lifespan which had independent biological replicates. Data distribution was assumed to be normal in bench experiments since parameters measured such as length, % proliferation and fertility are usually normal. However, this was not formally tested. In omics database analysis the data was normalized, the method of normalization for each dataset is listed in the Methods section.

## Data availability

All raw RNA sequencing data (bulk and single-cell) and Hi-C data, as well as processed datasets, can be found in the GEO database, accession numbers GSE248741 and GSE218971, respectively. Metabolomics data is available in **Supplementary Table S9**. All other data are available from the corresponding author upon request. All Supplementary Tables are available in Mendeley Data https://data.mendeley.com/datasets/ggys689v6x.

## Code availability

The code supporting the current study is available in the following GitHub repository: https://github.com/Harel-lab/germline-regulation-of-the-vertebrate-lifespan. The Hi-C code is available at https://gitlab.com/mcfrith/last/-/blob/main/doc/last-papers.rst.

## Acknowledgements

We thank Alon Zaslaver, Eran Meshorer, Anne Brunet, Yonanan Tzur, Nicholas E. Stroustrup, Mor Nitzan, and the Harel lab for stimulating discussion and feedback on the manuscript. We thank Ariel Velan, Ella Yanay, Ashayma Abu-tair, Yonatan Birenbaum, Fatma Idrees and Reem Barakat for help with killifish maintenance, Naomi Melamed-Book from the imaging facility (HUJI) and Sergey Malitsky and Maxim Itkin from the Life Sciences Core Facilities, Metabolic Profiling Unit (WIS). Supported by ERC StG #101078188 (I.H.), the Zuckerman Program (I.H.), Abisch-Frenkel Foundation 19/HU04 (I.H.), ISF 2178/19 (I.H.), Israeli Ministry of Science 3-17631 (I.H.), 3-16872 (I.H.), the Moore Foundation GBMF9341 (I.H.), BSF-NSF 2020611 (I.H.), the Israeli Ministry of Agriculture 12-16-0010 (I.H.), the Levi Eshkol scholarship of the Israeli Ministry of Science (E.M.), the Czech Science Foundation (#22-01781O), and the Ministry of Education, Youth and Sports of the Czech Republic (#CZ.02.1.01/0.0/0.0/16_025/0007370) (R.F.), the Chan Zuckerberg Initiative 2017-174468 and 2018-182817 (W.J.G.), and NIH grant numbers P50HG007735, UM1HG009442, 1UM1HG009436 (W.J.G.).

## Author Contributions

E.M., T.A., and I.H. designed the study. E.M. and R.F. performed experiments, E.M. generated the HPG and *dnd1* mutant killifish lines. X.S. and E.M. prepared the single-cell RNA libraries under the supervision of O.R. and I.H., respectively. T.A. designed and performed the analysis of single-cell RNA sequencing under the supervision of I.H.. A.S. performed worm lifespan experiments under the supervision of E.C.. T. A. and E. M. performed statistical analyses, with help from S.S. and D.M.Z.. A.O.G., G.K.M., and W.J.G. performed and analyzed the Hi-C data. T.A., E.M., and I.H. wrote the manuscript. All authors commented on the manuscript.

---

**Extended Data Figure 1:**
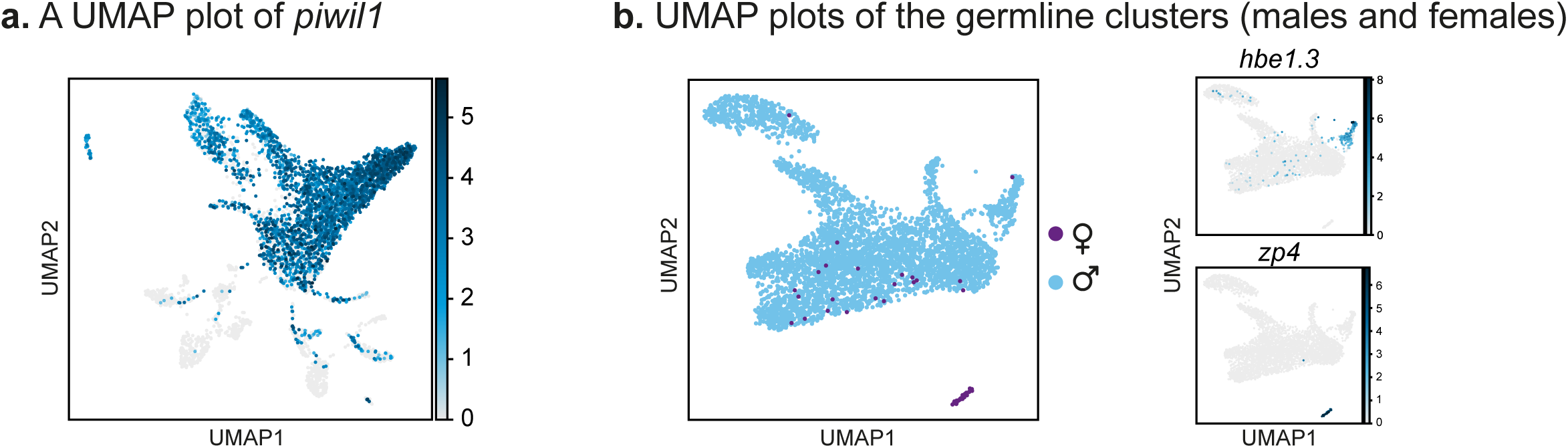
Filtering germ cell clusters. (**a**) Gene expression UMAP plot of *piwil1,* a germ cell marker gene. (**b**) Sub-clustering of the germ cells, performed according to Figure 1c, color-coded by sex (left). Within this cluster, a marker of erythroid cells (top right) and previtellogenic oocytes (bottom right) is also indicated. A full list of erythroid marker genes can be found in **Supplementary Table S3**.

**Extended Data Figure 2:**
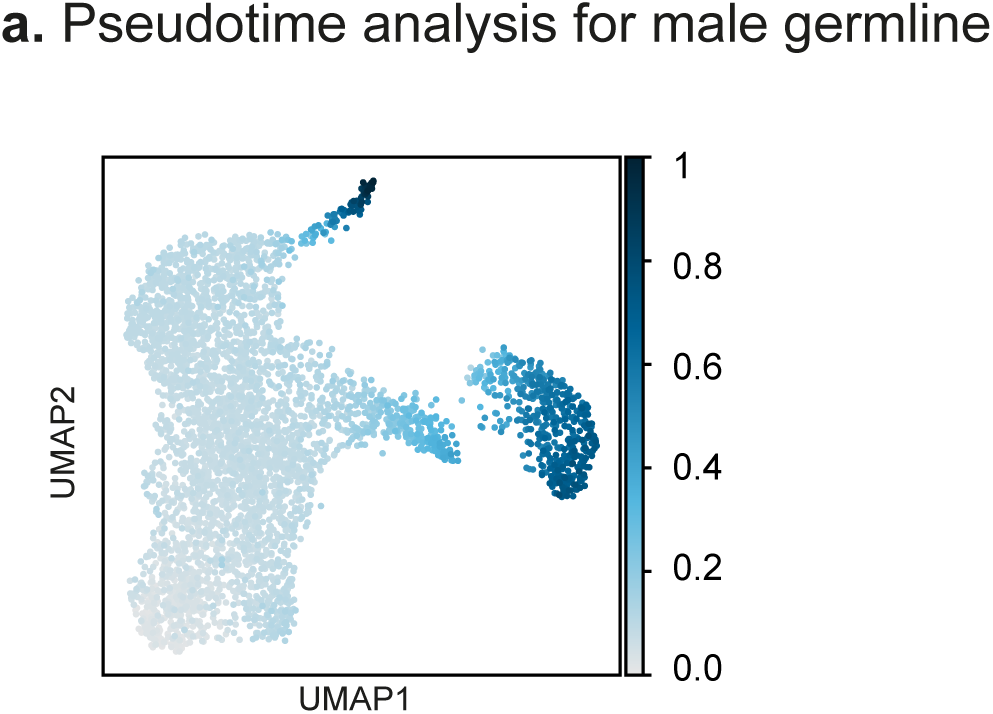
Pseudotime of germ cell clusters. (**a**) Sub-clustering of the male germ cells excluding the female germ cells and erythroid cells. Cells are color-coded and labeled pseudo-time.

**Extended Data Figure 3:**
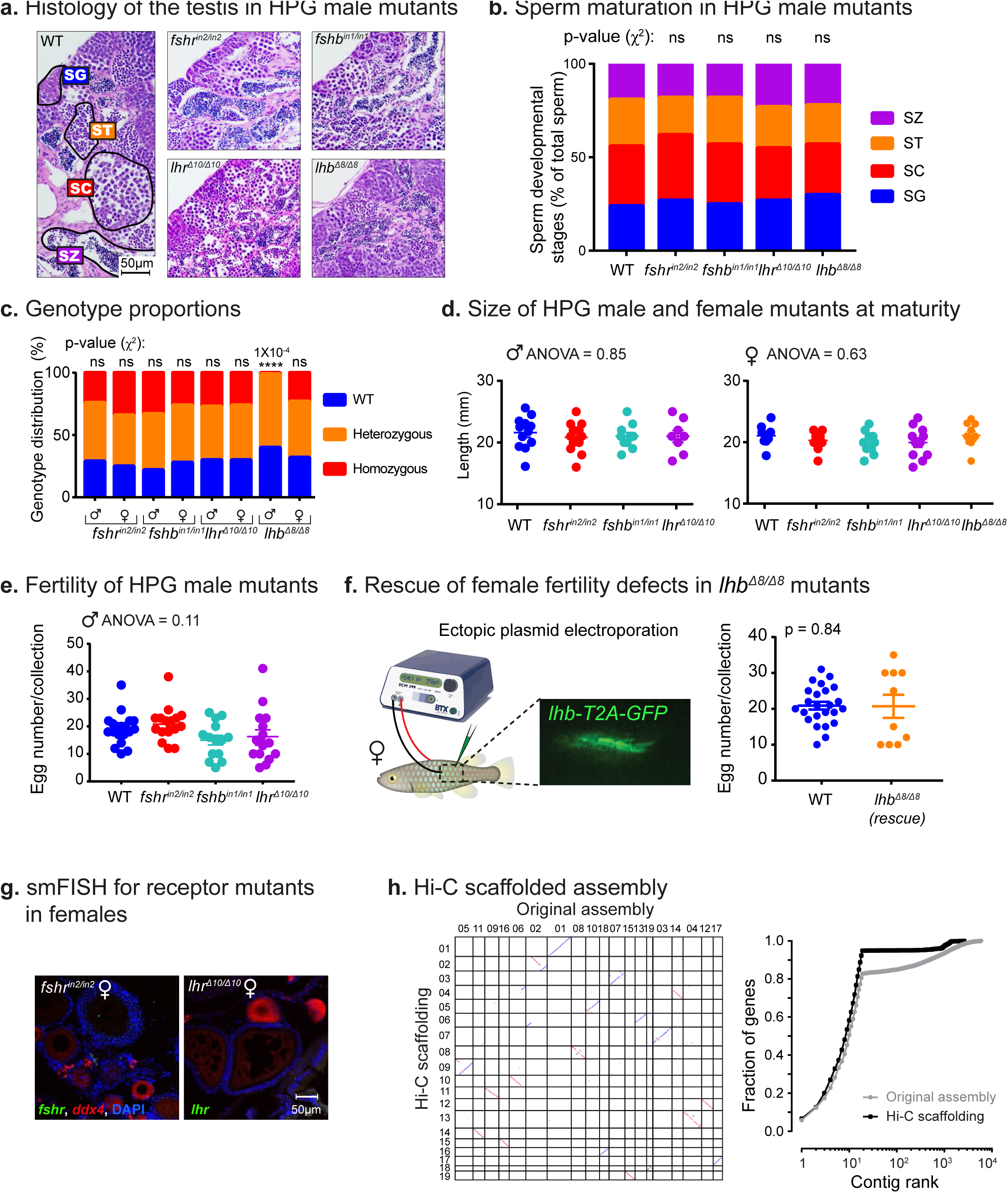
Perturbation and rescue of the reproductive axis in killifish. (**a**) Histological sections from one-month-old males. n ≥ 4 individuals from each genotype (except for *lhb*, which was sex-linked, n=1). Scale bar: 50 µm. Sperm developmental stages according to^102^: SG: spermatogonia; SC: spermatocytes; ST: spermatids; SZ: spermatozoa. (**b**) Quantification of sperm maturation, examples in (**a**). Data presented as proportion of each developmental stage. n ≥ 4 individuals for each experimental group (except for *lhb*). Significance was measured by a two-sided χ^2^ test with the WT proportion as the expected model. (**c**) Distribution of genotype progeny from heterozygous pairs (stratified by sex). n = 30-130 individuals, per genotype/sex. Significance was measured by a two-sided χ^2^ test with Mendelian proportions (25:50:25) as the expected model and FDR correction. (**d**) Quantification of somatic growth by calculating the standard length of one-month-old males (left) and females (right) of the indicated genotypes. n =8-12 individuals for all experimental groups. Error bars indicate meanLJ±LJSEM. Significance was calculated using one-way ANOVA. (**e**) Quantification of male fertility. Each dot represents the number of eggs per breeding pair, per week of egg collection (except for *lhb*, which was sex-linked). n=4-6 pairs for each group, over 4 collections. Error bars indicate meanLJ±LJSEM with individual points. Significance was calculated using one-way ANOVA. (**f**) Schematic illustration (left). Representative of n=3 females. Quantification of fertility output (right) in *lhb*^Δ8^*^/^*^Δ8^ mutant females following plasmid electroporation. Each dot represents the number of eggs per breeding pair per week. n = 3-6 pairs over 4 collections. Error bars show meanLJ±LJSEM. Significance was calculated using a two-sided Student’s t-test. (**g**) smFISH in the ovaries of the indicated genetic models. Marker for immature germ-cells (*ddx4*) in red, and for *fshr* and *lhr* in green. Representative of n ≥ 6 individuals. Scale bar: 50 µm. (**h**) Oxford Grid plots (left) showing correspondence between the Hi-C linkage groups (Y-axis) and previously predicted chromosomes (X-axis). Positive strand in blue, and negative strand matches in red. Cumulative fraction of genes (right) located on Hi-C pseudochromosomes and on contigs in the original assembly (gray). In the Hi-C scaffolded assembly (black), most small contigs containing genes are placed into the 19 main pseudochromosomes.

**Extended Data Figure 4:**
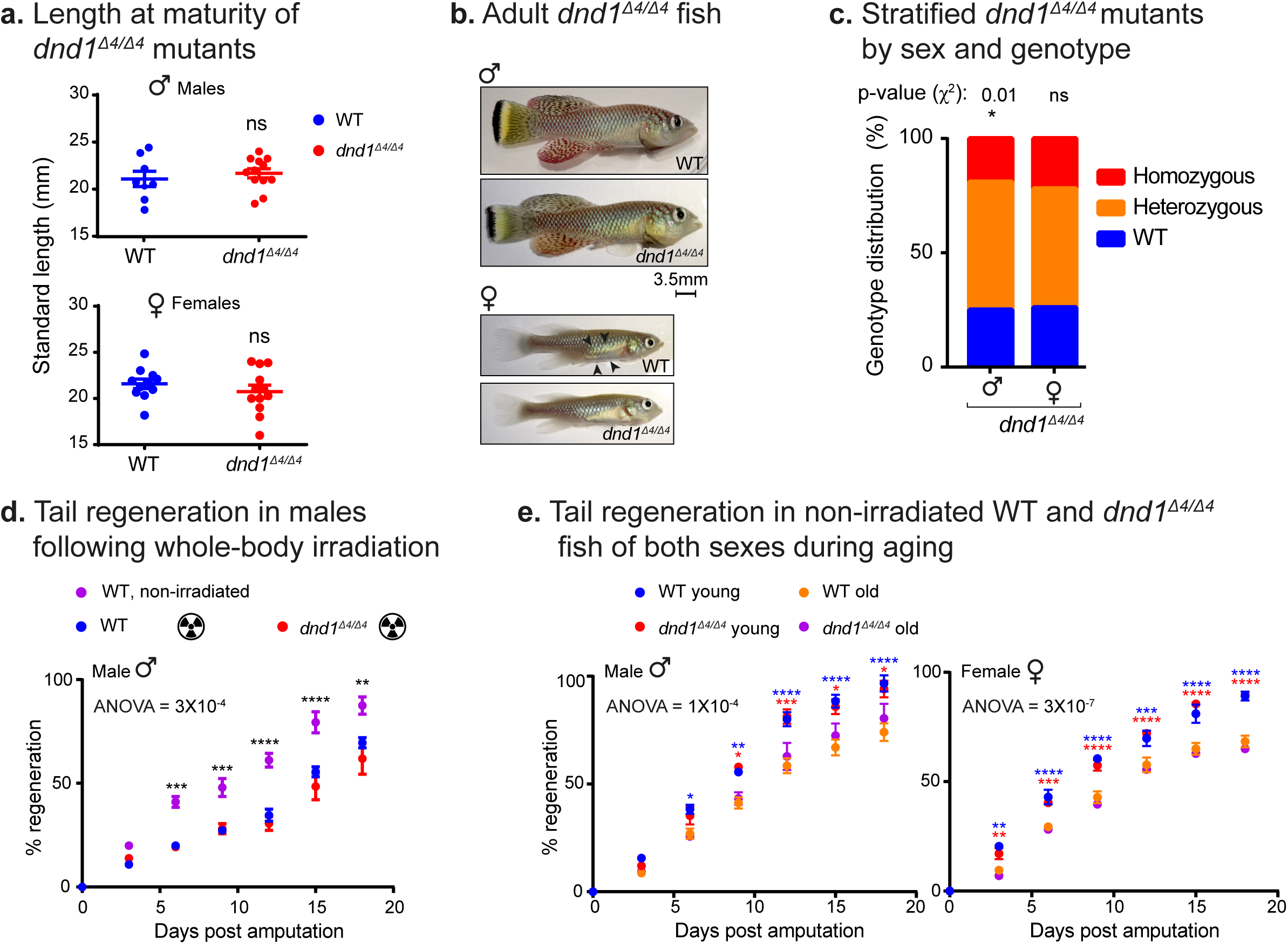
Physiological characterization germline-depleted fish. (**a**) Quantification of somatic growth (standard length) of WT or *dnd1*^Δ4^*^/^*^Δ4^ one-month-old mutants: males (top) and females (bottom), n = 8-12 individuals for all experimental groups. Error bars show meanLJ±LJSEM. Significance was measured using a two-sided Student’s t-test. (**b**) Representative images of adult (2.5-month-old) males (top) and females (bottom) WT or *dnd1*^Δ4^*^/^*^Δ4^ mutant fish. Black arrowheads highlight the presence of eggs in the female representative of n ≥4 individuals. Scale bar: 3.5 mm. (**c**) Distribution of genotype progeny from *dnd1*^Δ4^*^/+^* heterozygous pairs (stratified by sex). Significance was measured by two-sided χ^2^ test with Mendelian proportions (25:50:25) as the expected model and FDR correction. n ≥ 500 fish for each group and p-values are indicated. (**d**) Tail-fin regeneration following whole-body irradiation in males. Genotypes and treatments are indicated (n = 4 individuals for non-irradiated WT and 6 individuals for WT and *dnd1*). The length of the outgrowth was calculated as the percentage of the original fin size (before the amputation). Error bars indicate meanLJ±LJSEM. Significance was calculated using repeated-measures two-way ANOVA with a Dunnet post-hoc compared to the irradiated WT, and p-values are indicated. (**e**) Tail-fin regeneration during aging, in males (left) and females (right). Fish sex, genotypes, and age are indicated (n = 5-7 individuals for all experimental groups). Error bars indicate meanLJ± SEM. Significance was calculated using repeated measures two-way ANOVA with a Tukey post-hoc, and p-values of the comparison between young and old fish of each genotype are indicated by color-coding.

**Extended Data Figure 5:**
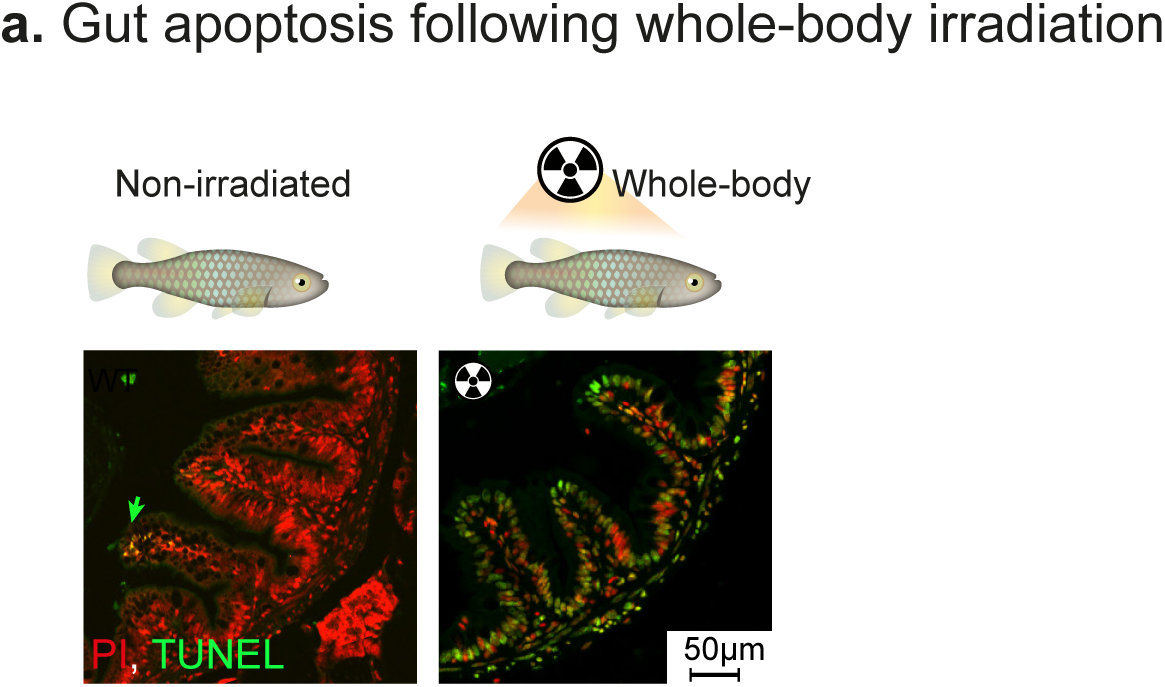
Calibration of DNA damage detection. (**a**) Representative images of apoptosis detected by the TUNEL assay (green and green arrows) and DNA (PI, in red). Assay was performed in gut sections of non-irradiated (left) or irradiated fish (right). n=6 individuals in each experimental group. Scale bar: 50 µm.

**Extended Data Figure 6:**
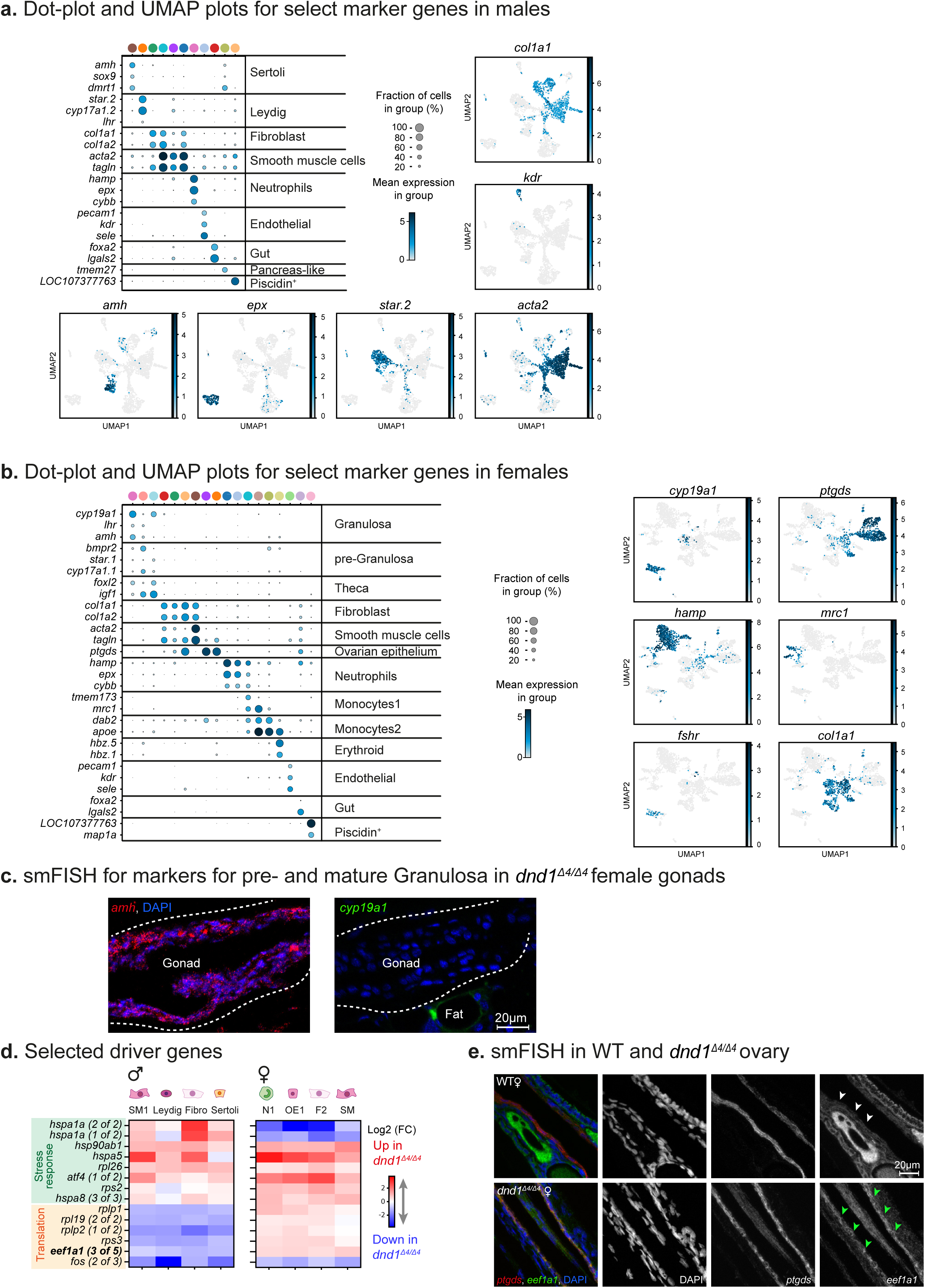
Characterization of cell type markers and differential gene expression in *dnd1*^Δ4^*^/^*^Δ4^ mutant gonads. (**a-b**) Dot-plot (top left, **a**, left, **b**) and UMAPs presenting the expression of select marker genes for males (**a**) and females (**b**). Dot-plot clusters are color-coded according to Figure 6a, b (for males and females separately). Note that most of the markers overlap with Figure 1c. A full list of markers can be found in **Supplementary Tables S5, S6**. (**c**) smFISH in ovaries for selected markers in *dnd1*^Δ4^*^/^*^Δ4^ mutants. *Amh* (red), a marker for Pre- and mature Granulosa cells, and *cyp19a1* (green), a specific marker for mature Granulosa. Note that *cyp19a1* is expressed in the adjacent adipose tissue. Representative of n ≥ 4 individuals. Scale bar: 20 µm. (**d**) Log2 fold-change heatmap represents the gene expression ratio between WT and *dnd1*^Δ4^*^/^*^Δ4^ fish of the indicated cell-types. The genes were selected as they were similarly altered in several cell types. (**e**) smFISH in WT fish and *dnd1*^Δ4^*^/^*^Δ4^ ovaries. *Eef1a1,* a marker for translation initiation (green), and *ptgds*, a marker for ovarian epithelium (in red). Representative of n ≥ 4 individuals. Scale bar: 20 µm.

**Extended Data Figure 7:**
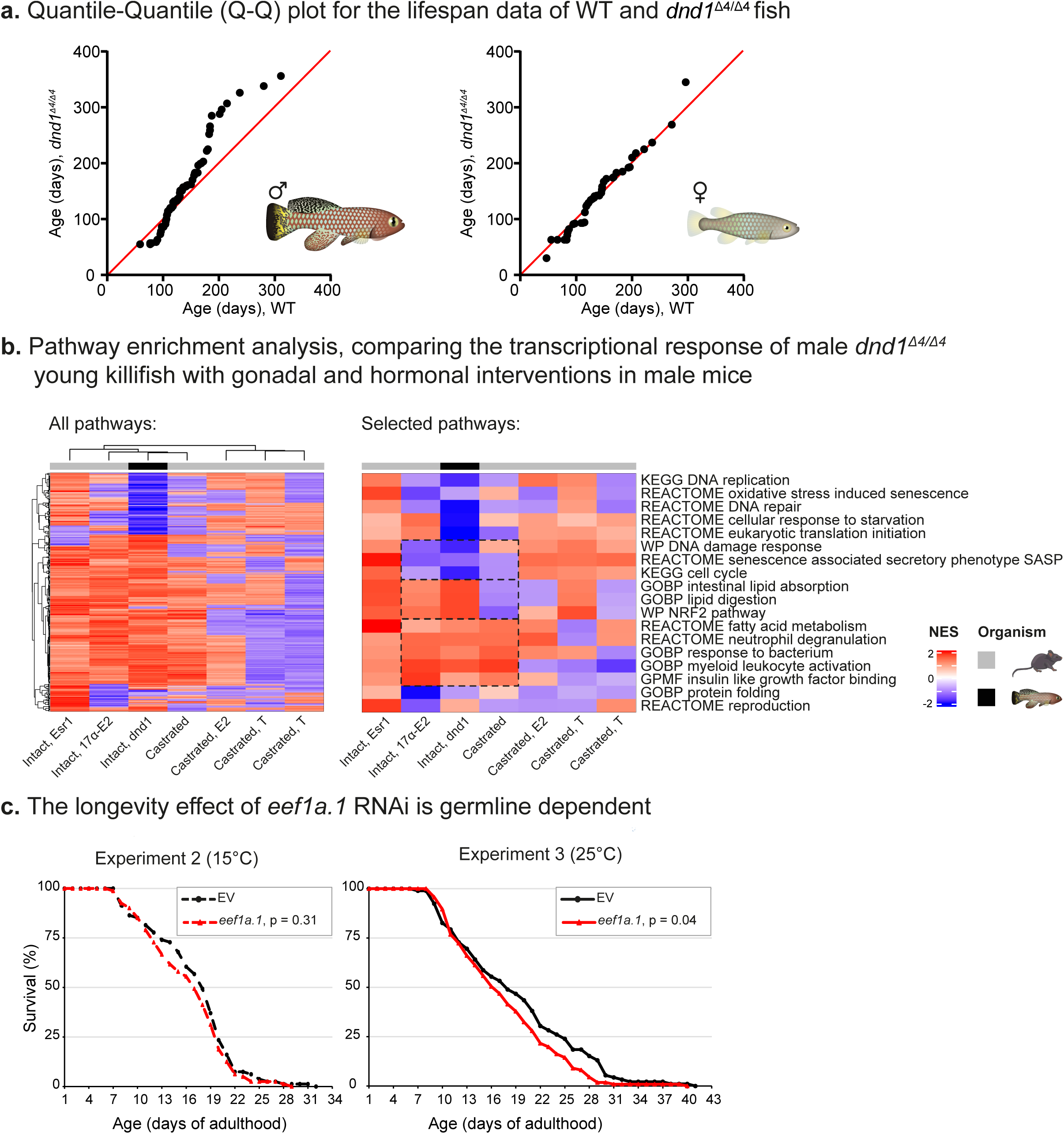
Germline depletion extends male maximal lifespan. (**a**) Quantile-Quantile (Q-Q) plots for the survival data of WT and *dnd1*^Δ4^*^/^*^Δ4^ fish (shown in Figure 7b), assessed separately for males (left) and females (right). (**b**) Heatmap for normalized enrichment scores (NESs) in the male liver, comparing the response to gonadal and hormonal treatments in mice with germline-depleted young fish (left, selected pathways highlighted on the right). A full list of pathways can be found in **Supplementary Table S8**. Esr1: estrogen receptor KO; E2: estrogen treatment; T: testosterone treatment. (**c**) Lifespan of *C. elegans* from the *glp-1*(e2144) mutants, grown either at the permissive 15°C (fertile, left) or restrictive 25°C (germline depleted, right), and fed with either EV or *eef1a1* RNAi. P-values were calculated according to log-rank. Worm numbers and raw data can be found **Supplementary Table S4**.

**Extended Data Figure 8:**
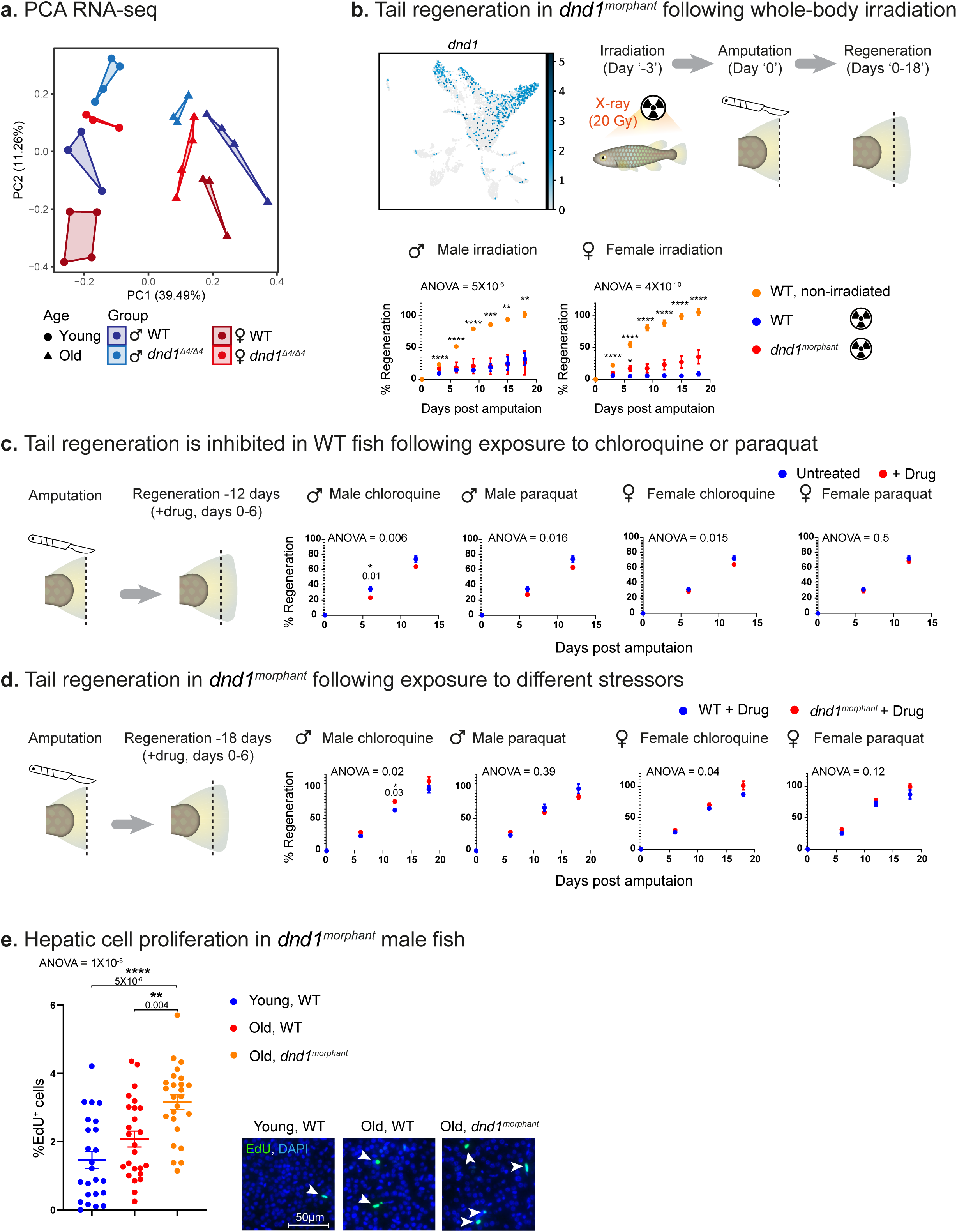
Germline depletion enhances regeneration under several stressors. (**a**) PCA for hepatic transcript levels. Groups include male (blue) or female (red), WT (dark shades) or *dnd1*^Δ4^*^/^*^Δ4^ (light shades), either young (circles) or old (triangles). n=3-4 samples per condition, and each symbol represents an individual fish. (**b**) Top, left: gene expression UMAP plot of *dnd1* gene. Right and bottom: tail-fin regeneration of WT and *dnd1^morphant^*fish following whole-body irradiation. n = 4 individuals for WT and *dnd1^morphant^* males, 7 for WT and *dnd1^morphant^* females and 10 for non-irradiated males and females. Error bars indicate meanLJ± SEM. Significance was calculated using repeated-measures two-way ANOVA with a Dunnet post-hoc, compared to WT irradiated fish, and p-values are indicated. (**c**) Tail-fin regeneration of WT fish following treatment by chloroquine or paraquat. Treatments and sexes are indicated. n = 5 individuals for untreated females, 11 for chloroquine treated females, 12 for untreated and chloroquine treated males, and paraquat treated females and 15 for paraquat treated males. The same untreated fish were used as control for both treatments. Error bars indicate meanLJ± SEM. Significance was calculated using repeated-measures two-way ANOVA with a Sidak post-hoc, and p-values are indicated. (**d**) Tail-fin regeneration of WT and *dnd1^morphant^* fish following treatment by chloroquine or paraquat. Treatments and sexes are indicated. n = 5 individuals for chloroquine treated *dnd1^morphant^* males, 7 for paraquat treated *dnd1^morphant^* males and chloroquine treated *dnd1^morphant^* females, 8 for chloroquine treated WT and paraquat treated WT and *dnd1^morphant^* females, 9 for paraquat treated WT males and 10 for chloroquine treated WT males. Error bars indicate meanLJ± SEM. Significance was calculated using repeated-measures two-way ANOVA with a Sidak post-hoc, and p-values are indicated. (**e**) Right: Representative images of proliferation detected in the livers of male fish of the indicated experimental groups, following a 3-day treatment of EdU (green, see **Methods**) with DAPI (blue) for detecting nuclei. Representative of n =5-7 individuals per group. Scale bar: 50 μm. Left: Quantification of the percentage of proliferating cells. Error bars indicate meanLJ± SEM. Significance was calculated using one-way ANOVA with a Tukey post-hoc, and p-values are indicated.

